# A model of flexible motor sequencing through thalamic control of cortical dynamics

**DOI:** 10.1101/2019.12.17.880153

**Authors:** Laureline Logiaco, L.F. Abbott, Sean Escola

## Abstract

The mechanisms by which neural circuits generate an extensible library of motor motifs and flexibly string them into arbitrary sequences are unclear. We developed a model in which inhibitory basal ganglia output neurons project to thalamic units that are themselves bidirectionally connected to a recurrent cortical network. During movement sequences, electrophysiological recordings of basal ganglia output neurons show sustained activity patterns that switch at the boundaries between motifs. Thus, we model these inhibitory patterns as silencing some thalamic neurons while leaving others disinhibited and free to interact with cortex during specific motifs. We show that a small number of disinhibited thalamic neurons can control cortical dynamics to generate specific motor output in a noise robust way. If the thalamic units associated with each motif are segregated, many motor outputs can be learned without interference and then combined in arbitrary orders for the flexible production of long and complex motor sequences.

## 1 Introduction

Animals have the remarkable ability of flexibly performing long and complex sequences of movements [42, 13, 52, 28]. In humans, dance provides an illustration of this ability. A long dance can be decomposed into a sequence of short stereotyped moves or motifs. These motifs form a library that can be flexibly combined by experienced dancers to create novel sequences – new dances – with minimal additional training. Furthermore, new motifs can be learned to extend the library without interfering with previously acquired dance moves. These phenomena raise important questions regarding sequence generation in the mammalian motor system that have not yet been addressed with computational models. First, how is flexibility achieved? This is an open issue – artificial neural networks struggle to generalize to situations not encountered during training [2, 14, 37]. A motor analog would be a previously unexperienced transition between a pair of known motifs [31]. Second, how can a motif library be extended? This is another problem with a parallel in artificial networks known as catastrophic forgetting; the overwriting of prior knowledge during new learning [20, 43]. More generally, how can a high level sequencing command, instructing which motifs to execute and in which order, be efficiently communicated to a neural network dedicated to the dynamic elaboration of the corresponding motor program?

The anatomy and physiology of the motor system may provide clues to and constraints on the answers to these questions. We focus on the recurrent system comprising motor cortex, the basal ganglia, and thalamus for which a significant body of evidence supports a role in the learning and execution of skilled motor sequences [13, 45, 25, 48, 39, 24]. Specifically, the motor cortex appears to function as a dynamic pattern generator that produces neural activities needed to execute the muscle contractions associated with individual motifs [6, 48]. The basal ganglia, on the other hand, have been linked to computations needed for arranging motifs into sequences. The striatum may control sequence structure [13] while the output nuclei – the internal capsule of the globus pallidus (GPi) and substantia nigra par reticulata (SNr) – generate sustained firing patterns that are specific to particular motifs [25]. During sequence generation, the motor thalamus is typically considered to function as a relay, receiving strong inhibitory input from the basal ganglia [10, 9] and projecting to cortex [49]. However, the motor thalamus also receives feedback from motor cortex [18]. Thus, in addition to the conventional loop from cortex through the basal ganglia and thalamus and back to cortex, there is a second loop directly between cortex and thalamus. While the former has long been studied [1], only recently has the functional importance of the second loop been characterized [44, 46, 17]. Lacking, however, is a model of how motor function is supported by the interaction between these two loops.

We have developed a model that incorporates thalamocortical loops controlled by basal ganglia outputs that can flexibly and extensibly generate sequences (Fig. 1a). The model includes bidirectionally connected motor cortex and thalamus and inhibitory connections from basal ganglia to the thalamus. The model generates muscle-like activity patterns through an output unit formed from a weighted sum of cortical activities. The basal ganglia inputs can silence thalamic neurons but, otherwise, the thalamic and cortical regions are modeled as linear, making the full model equivalent to a switching linear system [30]. This allows for a complete analysis that reveals the relationship between system output and the synaptic weights of the cortex–thalamus circuit. We find that the dynamics needed to execute a specific motif in a noise-robust manner can be generated solely by adjusting the synapses between cortex and motif-specific thalamic units; all synapses within cortex and to the output unit are unchanged by learning.

**Figure 1:**
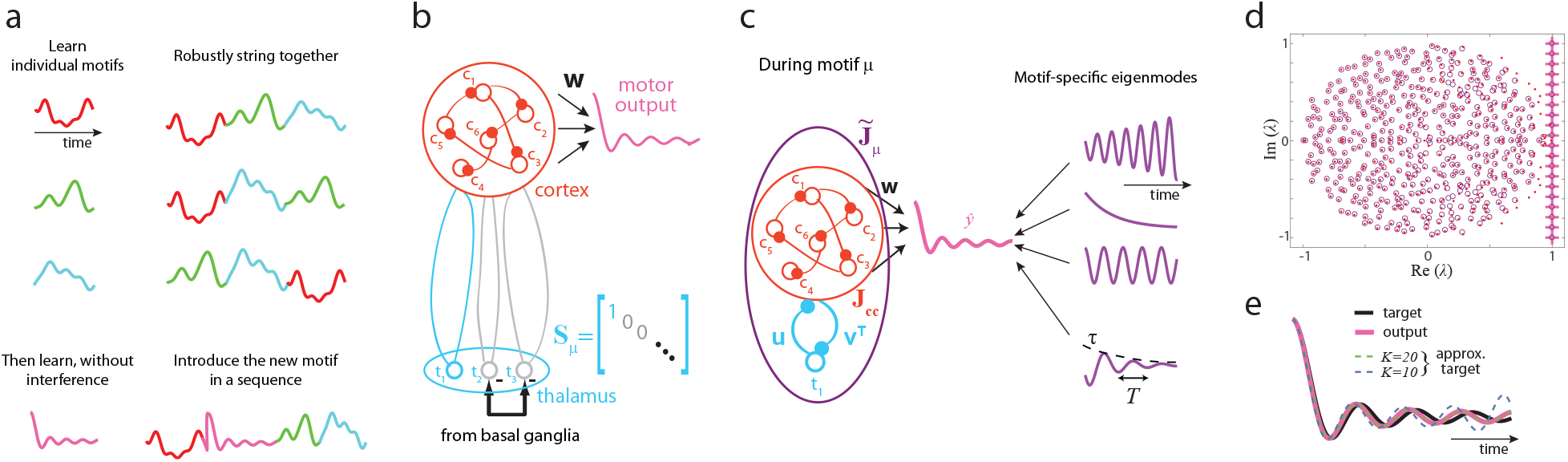
Motif sequencing and the cortex–thalamus circuit. **a**) Flexible and extendable motor sequencing. Our goals are to execute motor motifs from a library (top left) in sequences of arbitrary order (top right) without training all possible transitions, and to learn a new motif (bottom left) without interfering with the existing library for incorporation into novel sequences (bottom right). **b**) Cortex–thalamus model schematic. Cortical network in red, is reciprocally connected to non-recurrent thalamic units that receives inhibitory input from the basal ganglia. During the execution of a motif, basal ganglia activity inhibits selected thalamic units (e.g., *t*_2_ and *t*_3_) and leaves others active (e.g., *t*_1_), as captured in the diagonal selection matrix **S**_*μ*_ for motif *μ* (Eq. 3). The cortex–thalamus circuit generates motor output 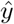 through readout weights **w**. **c**) Motif-specific dynamics. The effective connectivity matrix 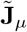 defines the dynamics of motif *μ*. Modes of the cortex–thalamus circuit (purple traces) are summed to produce a desired motor output (pink trace). The real and complex parts of the eigenvalues of 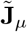 define respectively the exponential decay times *τ* and oscillation periods *T* of each mode. **d**) Eigenvalue control during a motif. The target eigenvalues (+ symbols) are included in the spectrum of the cortex–thalamus circuit resulting from imposing the relationship between the corticothalamic and thalamocortical synaptic weights given by Eq. 6 (for *K* = 20, purple circles). The spectrum of the purely cortical network is shown for comparison (red dots). *N* = 500. **e**) Approximating a target motor motif (black curve) by a weighted sum of *K* complex exponential modes of the cortex–thalamus circuit with *K* = 10 (green dashed curve) or *K* = 20 (blue dashed curve). The pink curve is the system output when the eigenvalues are given by the purple circles in **d**.

To generate an arbitrary sequence, the basal ganglia switches between inhibiting different subsets of thalamic units to execute different motifs. This framework ascribes to the basal ganglia the role of sequence selection and to the thalamocortical loop, in conjunction with motor cortex, the role of motif execution. The model predicts that experimental manipulations of the basal ganglia or cortical inputs to motor thalamus should differentially affect selection and execution, respectively. Finally, our framework suggests that the cortex–thalamus architecture is well-suited to flexibly control prolonged and complex sequential motor behavior.

## 2 Results

Our aim is to develop an understanding of how cortical and subcortical motor areas cooperate to implement flexible motor sequencing. In particular we wish to gain insight into how the anatomy of the motor system supports these computations. To do this, we constructed a model that is faithful to the general anatomy but simple enough to be analytically tractable (Fig. 1b, and see Supplementary Table 5.2.1 for a complete list of the assumptions in our model). On the basis of previous experimental and theoretical work [48, 6, 50], we assume that motor cortex dynamics generates patterns of activity that form a basis for generating the muscle activity associated with movement. We use a standard ‘firing-rate’ description in which the cortical activity, described by a vector **c**, interacts with thalamic activity **t**, and generates motor output 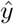 through the equations

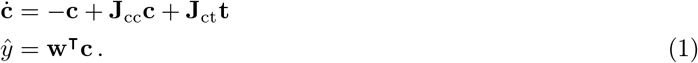

The components of the vectors **c** and **t** represent the activities of ‘units’, corresponding to linear combinations of neural activities. **J**_cc_ and **J**_ct_ are intracortical and thalamocortical synaptic weight matrices, and **w** is the vector of output weights used to generate the signal 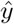. The thalamic activity **t** could be given by an equation similar to that for **c** but, instead, we assume that the dynamics in thalamus are more rapid than in cortex and approximate the thalamic response to cortical activity as instantaneous and proportional to the product of a corticothalamic weight matrix **J**_tc_ and the cortical activity **c**. (The results we present below are qualitatively unchanged if we relax these assumptions; see Supplementary Sec. 5.1.5.)

Thus far, the model we have described is linear, but we introduce a critical nonlinearity in the model by allowing thalamic units to be switched ‘off’ and ‘on’. The inputs from GPi/SNr (i.e., the output nuclei of the basal ganglia) to motor thalamus are strong and inhibitory; and the activity patterns in GPi/SNr are sustained during motif execution, with switches predominantly at the transitions between motifs [25]. In our model, we assume that the synapses from basal ganglia to thalamus are strong enough that, when active, they completely inhibit their thalamic targets (though we can also relax this assumption as described in Supplementary Sec. 5.1.4). This is compatible with the classic view that motor thalamus is in an inhibited state by default and is disinhibited by targeted removal of inhibitory input from the basal ganglia, which gates movement [9, 10, 27], but see [47]. During a particular motif, the thalamic units that are not inhibited by the basal ganglia are assumed to be in the linear regime, while those that receive basal ganglia input are silent. With these assumptions, we can express the activity of the thalamic units during motif *μ* as

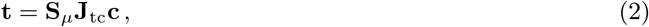

where **S**_*μ*_ is a diagonal selection matrix whose only nonzero elements are 1s at locations corresponding to thalamic units that are active during motif *μ*.

Combining Eqs. 1 and 2, we obtain a closed equation for the cortical activity during motif *μ*,

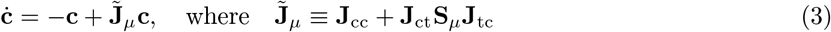

Thus, during a particular motif, cortical dynamics are governed by the *effective* connectivity matrix 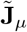 given by the sum of the fixed corticocortical connectivity **J**_cc_ and a motif-specific corticothalamocortical perturbation matrix **J**_ct_**S**_*μ*_**J**_tc_. In the following sections, we study this model and show that selective thalamic silencing by the basal ganglia can flexibly remap cortical dynamics and control the output 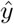.

### 2.1 Executing a target motif

The time-dependent cortical activity for motif *μ* generated by Eq. 3 is completely determined by a set of initial conditions, **c**(0) and by the effective connection matrix 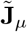. Specifically, if 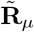 and 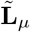 are matrices of the right and left eigenvectors of 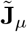 and 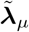 is the vector of its eigenvalues, all associated with motif *μ*, then

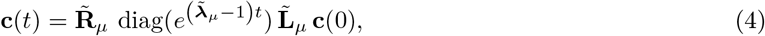

where we define *e*^**x**^ as the vector whose elements are the exponentials of the elements of **x**. Note that, for notational simplicity, time *t* in **c**(*t*) is the time elapsed since the beginning of motif *μ*, not the time since the beginning of the entire sequence of motifs.

From this, if we want to generate an output through a weighted sum of the complex exponential modes of 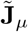 – i.e., 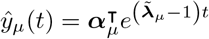 for some vector of coefficients ***α***_*μ*_ (Fig. 1c) – we need to set

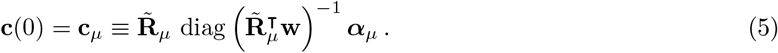

In practice, we compose 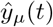 using only *K* modes, with *K ≪ N*, which means that the remaining components of ***α***_*μ*_ are zero. 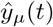 itself is an approximation to some desired output motif *y_μ_*(*t*), and when considering a variety of desired outputs, we find that 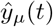 approximates *y_μ_*(*t*) faithfully for *K* ≲ 20 (i.e., ≲ 10 complex conjugate eigenmode pairs; Fig. 1e and see Supplementary Fig. S.2).

Executing a motif through Eqs. 4 and 5 requires that two things happen before the start of each motif. First, the eigenvalues described by the first *K* elements of 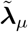, denoted by 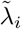 for *i* = 1, 2, …, *K*, must be set so that they produce oscillations and exponential decays that match the form of the desired output 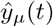. In the following section, we show how this can be done through selective silencing of thalamic units. Second, the initial condition of Eq. 5 must be established, and this must be done rapidly to avoid temporal gaps between motifs. In section 2.4, we show how this can be done through a combination of input to cortex and a different pattern of thalamic silencing. Since it is unreasonable for a biological system to perfectly achieve the initial condition given by Eq. 5, in section 2.3 we show that the thalamocortical and corticothalamic weights can be set such that even with noise in the initial conditions, motifs can be executed with high fidelity.

### 2.2 Thalamic silencing remaps cortical dynamics through thalamocortical loops

An hypothesis of our model is that all control of cortical dynamics arises from basal ganglia inhibition (silencing) of thalamic units. This requires that the activity of the large cortical network can be dramatically altered by a much smaller number of thalamic units. To test whether this is possible, we consider the minimal case in which a single thalamic unit is left free to interact with cortex during motif *μ*. In this case, the thalamic selection matrix **S**_*μ*_ is only non-zero for the entry referencing the active thalamic unit, and thus the corticothalamocortical perturbation **J**_ct_**S**_*μ*_**J**_tc_ reduces to a rank one matrix formed through the outer product of the *μ*^th^ column of the thalamocortical weights **J**_ct_ and the *μ*^th^ row of the corticothalamic weights **J**_tc_, which we denote as **u** and **v**^T^ respectively (dropping the *μ* index from **u** and **v** to streamline the notation; Fig. 1c). The effective connectivity then becomes 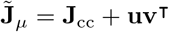.

To execute a particular motif, the connectivity vectors **v** and **u** must be chosen so that *K* of the eigenvalues of 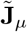 are set to the required values for motif *μ*. To do this, we make use of a relationship between these eigenvalues and **v** and **u** (see Methods section 4.1):

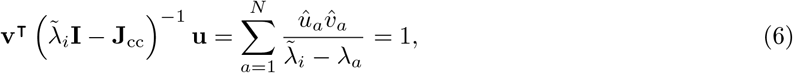

which is true for all *i*. In the second expression, 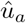 and 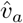 are the projections of **u** and **v** onto the *a*^th^ right and left eigenvectors of **J**_cc_, respectively, and *λ_a_* is the corresponding eigenvalue, for *a* = 1, 2, …, *N*.

By specifying *K* of the eigenvalues, Eq. 6 places *K* constraints on the 2*N* variables comprising **u** and **v**. If we define the vector **d** with elements 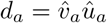 and the *K* × *N* matrix **P** with elements 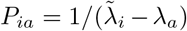, Eq. 6 can be rewritten as the linear system of equations 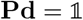 (where 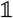 is a vector of all ones). Solutions to this system – and there are infinitely many because the system is underconstrained – give us **d**. Then, given either **u** or **v** (e.g., by choosing one of these connectivities randomly), the other can be calculated. Formally, this means that, starting with almost any cortical connectivity, we can obtain any desired spectrum of new eigenvalues – and therefore execute any motif we want – by choosing an appropriate thalamocortical loop. In practice, however, only under some conditions will (*i*) the corticothalamocortical perturbation weights be stably defined and have appropriate magnitude, and (*ii*) the cortex–thalamus circuit be robust to noise during motif execution. We discuss the first point here and the second in the following section.

The success of using Eq. 6 to control the eigenvalues of the cortex–thalamus circuit 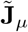 depends on the number of desired eigenvalues *K* and how far they fall outside of the spectrum of the initial corticocortical connectivity **J**_cc_. These determine to what extent the rows of **P** are correlated, and thus our ability to solve for thalamocortical and corticothalamic weights **u** and **v** that control the eigenvalues as desired. Luckily, as discussed in section 2.1, the motifs we consider can be executed using a small number of modes (i.e., *K ≪ N*), and we find that, in this case, setting the desired eigenvalues succeeds (Fig. 1d and Supplementary Fig. S.1) with corticothalamocortical weights of reasonable magnitude (Fig. 2c & f and Supplementary Fig. S.4). For this to work, how we choose to ‘invert’ **P** to solve for **d** – a non-unique operation – is important because certain solutions can stabilize the overall circuit dynamics (Supplementary Sec. 5.1.1). Furthermore, setting the desired eigenvalues successfully for each motif is facilitated by starting with a cortical circuit with rich dynamics, avoiding the need to change the existing cortical eigenvalues by large amounts. Specifically, we used a large random matrix **J**_cc_, which has eigenvalues distributed uniformly within the unit circle including some eigenvalues with real parts near one [15, 16]. These slow eigenmodes are a good initial basis for creating slow motor commands [22, 8].

**Figure 2:**
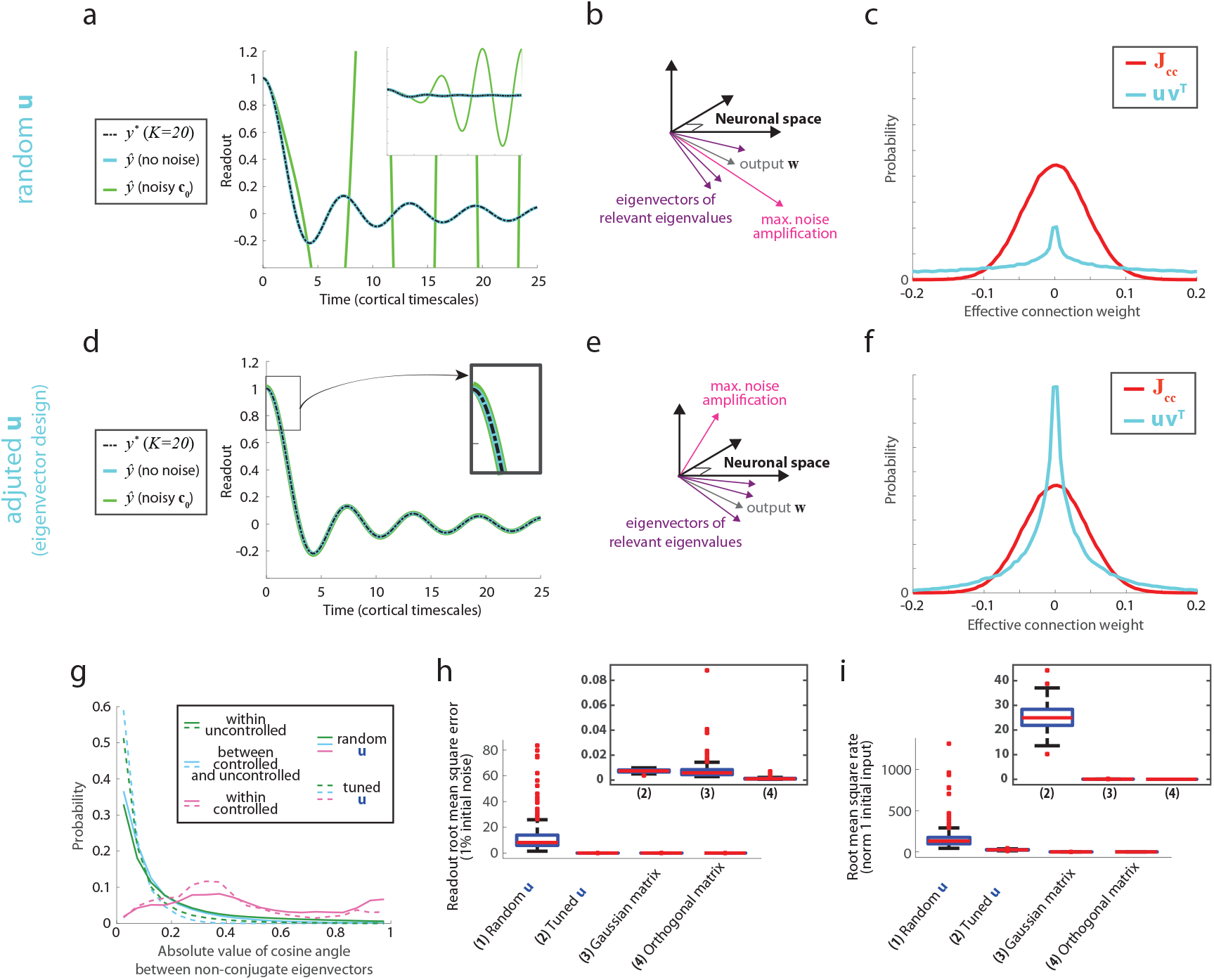
Noise robust motif execution. **a**) Target and system output in the presence and absence of noise for random **u**. With the initial activities **c** (0) set by Eq. 5 (i.e. **c**(0) = **c**_*μ*_), the output 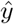 (cyan) matches the target (dashed black; the *K* = 20 curve in Fig. 1e). If noise is added to **c**(0), the output deviates wildly from the target (green and inset). Noise is Gaussian i.i.d. with standard deviation equal to 1% of the root mean squared norm of the activities during the motif in the absence of noise. **b**) Amplification of noise due to nonorthogonal eigenvectors. If the output weight vector and the direction of noise amplification are aligned, the output is highly noise sensitive. **c**) Distributions of cortical and loop weights for a random **u** sampled from a normal distribution. The effective corticothalamocortical weights in **uv**^T^ (standard deviation *σ* = 0.719) are more widely distributed than the corticocortical weights (*σ* = 0.045). **d–f**) Same as **b–d** but with the thalamocortical weights **u** optimized to minimize the effect of noise on the initial conditions. Noise has a negligible effect on the output (**d**) because the direction of noise amplification and the output vector are no longer aligned (**e**). Additionally, the distribution of the loop weights narrows (**f**, *σ* = 0.074). **g**) Distributions of the absolute values of the cosines between eigenvectors. Larger values indicate almost parallel eigenvector pairs (see Supplementary Sec. 5.1.2 for details). Larger correlations are observed when both eigenvectors correspond to controlled eigenvalues (magenta) than if only one (cyan) or neither (green) does. These correlations are similar whether **u** is random (solid) or optimized (dashed). **h**) Root mean squared error of the output in the presence of 1% noise. Compared to the case of random **u**(1), the noise-induced error is substantially diminished after optimization (2) and is on par with errors observed using matrices with the same eigenspectrum as 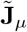 but with eigenvectors from random matrices (3) or orthogonal eigenvectors (4) (the latter yield explicitly normal dynamics, while the former support very limited non-normal amplification). See Supplementary Sec. 5.1.2 for details. **i**) Root mean squared norm of the activity vector **c**(*t*) when **c**(0) is a random vector of length one. Optimizing **u** reduces the length of the activity vector (1 vs. 2), but not to lengths observed with random or orthogonal eigenvectors (3 & 4), indicating that optimization does not eliminate nonnormal amplification. The data in **g**–**i** were generated from 50 random networks (i.e., random samples of **J**_cc_ and **w**) with five randomly sampled **u** per network; **h** & **i** used five matrices with random and orthogonal eigenvectors per network. *N* = 500 throughout.

In summary, thalamus can act as a controller of cortical dynamics, with even a single thalamic unit able to introduce multiple specified eigenmodes into the circuit dynamics, provided that: (*i*) the target eigenvalues are reasonably small in number (i.e., *K* is not of order *N*) and are not unreasonably far from the initial spectrum; (*ii*) the synaptic weights of the thalamocortical loops can be adjusted [38, 53, 23, 3]; and (*iii*) the loop weights can be moderately larger than the corticocortical weights.

### 2.3 Taming sensitivity to initial conditions

Even when we can successfully set the eigenvalues of the cortex–thalamus network with weights with reasonable amplitudes, undesirable properties of the eigenvectors of 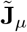 can cause problems for executing desired motifs. This problem comes about when eigenvectors are nonorthogonal – in particular, if some eigenvectors are almost parallel – and indeed we can show analytically that the process of controlling eigenvalues discussed in the previous section introduces near-parallelism between eigenvectors (see Fig. 2g and Supplementary Sec. 5.1.2). In this case, small fluctuations in the initial value of **c**(0) away from that specified by Eq. 5 can cause **c**(*t*) to deviate wildly from the desired motif trajectory. This is due to the well-known transient amplification phenomenon in linear networks with non-normal eigenvectors [12, 36, 21, and see Supplementary Sec. 5.1.2 and Supplementary Fig. S.3]. Without a reduction of this sensitivity to initial conditions, the system we propose is not viable because a small amount of noise can lead to catastrophic errors in the output (Fig. 2a).

Fortunately, we can minimize sensitivity to noise by taking advantage of the fact that the solution we obtain from Eq. 6 defines **d**, not the connectivity vectors **u** and **v**, leaving us with additional degrees of freedom. Thus far, we have chosen **u** randomly and then calculated the corresponding **v**. To stabilize against noise in the initial conditions **c**(0), we can instead choose a specific connectivity **u** as follows. First, we construct a cost function that quantifies the effect of noise in the initial conditions on the network output (Methods section 4.2). Then, we choose **u** to minimize this cost, with **v** determined from this **u** through Eq. 6.

After optimization, the system output is much less sensitive to noise (compare Figs. 2a & 2d) and has error magnitudes on par with those of reference matrices having little or no non-normal amplification (Fig. 2h). Interestingly, this reduction in noise sensitivity is not due to an elimination of non-normal amplication. In certain directions, small noise in the initial conditions still leads to deviations in network activity larger than for reference matrices (though reduced compared to the case of random **u**; Fig. 2i). These results indicate that optimizing the full thalamocortical loop stabilizes the output by making it orthogonal to the principal directions of noise amplification, while keeping it relatively aligned with the eigenvectors needed to construct the desired target motif (as schematized in Figs. 2b & 2e). In addition, optimizing for noise robustness makes the distribution of the weights of **uv**^T^ more closely match that of **J**_cc_ (Figs. 2c and 2f).

Thus, the small number of parameters associated with a single thalamocortical loop can be used to control the dynamics of the cortex–thalamus system such that complex output motifs can be executed in a noise robust manner.

### 2.4 Thalamocortical loops can prepare cortex to execute each motif

Executing a motif requires selecting a particular thalamocortical loop through basal ganglion input, as we have just described, and also establishing initial conditions that allow the resulting circuit to produce the desired output (Eq. 5). We now consider a mechanism by which cortical activity can be driven to the required motif-specific initial state. We assume that, following the completion of the previous motif but prior to the execution of motif *μ*, the motor system employs a preparatory period during which the cortex–thalamus network is driven by the dynamics 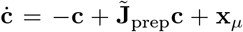, where **x**_*μ*_ is a motif-specific input. With these dynamics, the activity at steady-state will converge to **c**_*μ*_, the desired initial activity for motif *μ*, if

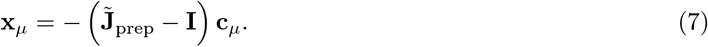

For this cortical input, the difference between the cortical state at the end of the previous motif and the desired initial state at the beginning of motif *μ*, denoted by *δ***c**, evolves according to 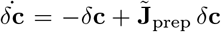. This formulation shows that the timescale of convergence to **c**_*μ*_ does not depend on either the previous motif or upcoming motif *μ* and thus that the same 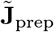 can implement transitions between all pairs of motifs provided that its eigenvalues have real part less than one. Furthermore, if the eigenvalues of 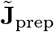 have real parts much smaller than one and if its eigenvectors do not create much non-normal amplification in the circuit, the transitions can be fast.

We have set the cortical matrix **J**_cc_ to have some eigenvalues with real parts near one to support the slow dynamics needed to execute motifs. Thus, these eigenvalues must be modified by the preparatory thalamocortical loop to assure rapid convergence to any desired initial condition and therefore successful and fast motif transitions. Because this requires modifying a considerable number of eigenvalues, it cannot be done stably using a single thalamic unit. To make transitions rapid, we consider a corticothalamocortical loop **J**_ct_**S**_prep_**J**_tc_, and optimize the relevant columns of **J**_ct_ and rows of **J**_tc_ to minimize the convergence time (Methods section 4.4). Using a thalamic population equal in size to 10% of the number of cortical units, convergence to steady-state occurs in just a few time constants of the cortical units (Fig. 3c). This occurs because the eigenspectrum has been shifted towards negative real values (Fig. 3d). Nevertheless, the synaptic weights of **J**_ct_**S**_prep_**J**_tc_ after optimization are reasonable in magnitude (Fig. 3e and see Methods section 4.4). Increasing the size of the thalamic population used leads to faster convergence times (Fig. 3f).

**Figure 3:**
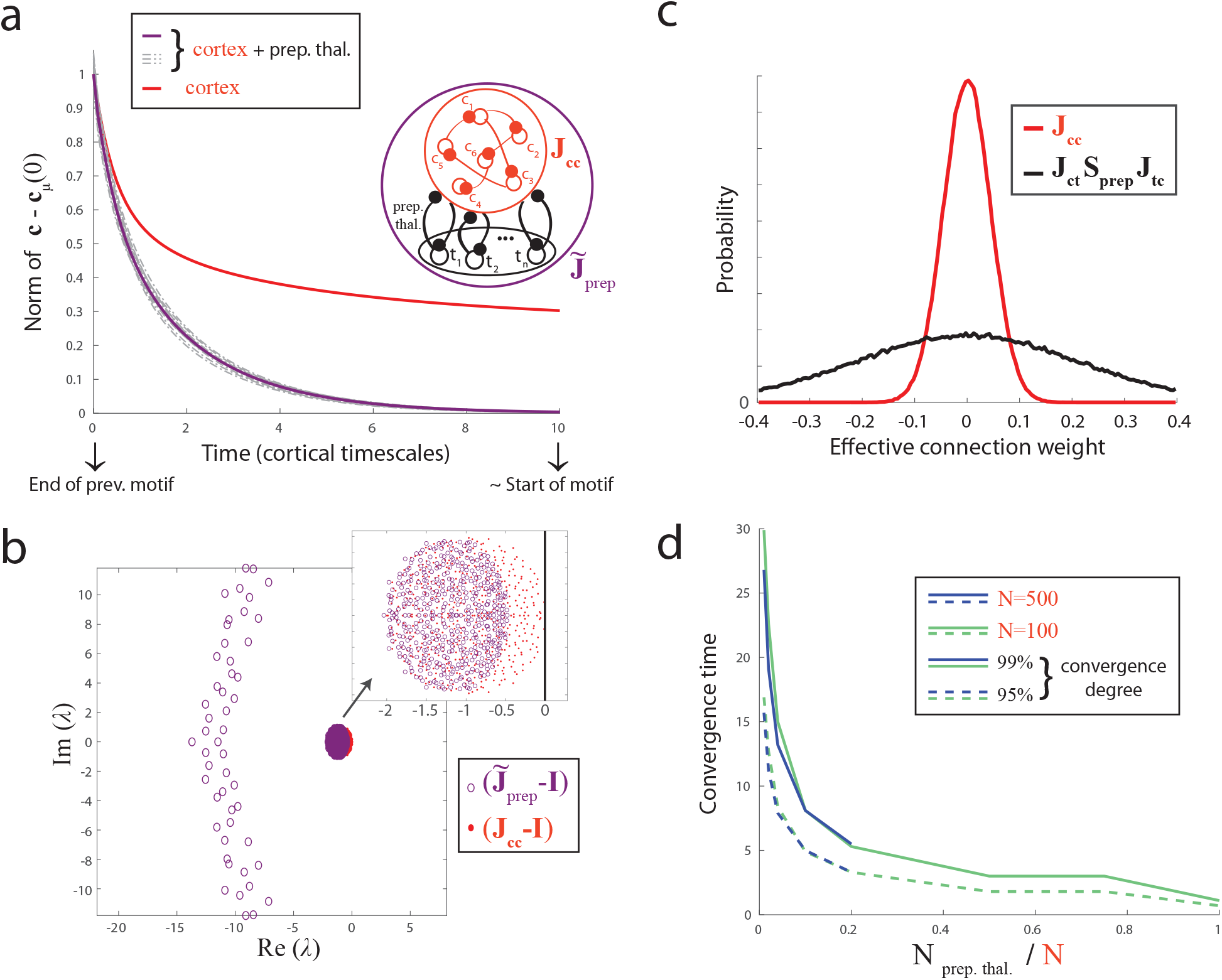
Facilitation of motif transitions by a preparatory thalamocortical network. **a**) The decay of the magnitude of *δ***c** for optimized preparatory networks (solid purple: average; dashed purple: individual trials) and for the purely cortical network (red, average). Optimizing the corticothalamocortical connections greatly speeds up the decay. With a thalamic population equal in size to 10% of the number of cortical units, convergence occurs in less than 10 timescales the time constant of the cortical units (defined as 1). **b**) Eigenvalues of the optimized preparatory cortex–thalamus network compared to those of the purely cortical network. **c**) Distributions of the synaptic weights of the corticothalamocortical preparatory weight matrix (black) and of the isolated corticocortical weights (red) (also see Supplementary Fig. S.4). **d**) Time for *δ***c** to decrease by either 95% (dashed) or 99% (solid) of its initial value as a function of the number thalamic units used in the preparatory network relative to the number of cortical units. For a fixed proportion of thalamic units, the decay time is similar between different numbers of cortical units (*N* = 500 in blue and *N* = 100 in green). *N* = 500 for **a**–**c**.

The cortex–thalamus preparatory network is active only during brief transitions between motifs. Some basal ganglia output neurons have been shown to fire transiently at the boundaries between sequence elements [24, 25], supporting the hypothesis that motor preparation is mediated by basal ganglia to thalamus interactions.

### 2.5 Switching between thalamocortical loops generates flexible, robust motor sequences

By alternating between loops that execute specific motifs and are robust to noise (as in Fig. 2d) and a fast preparatory loop (Fig. 3a), the cortex–thalamus circuit can generate arbitrary motif sequences (Fig. 4). Our model assumes that the basal ganglia select the active loop and thus the effective cortical dynamics through their strong inhibitory outputs to the thalamus (Fig. 1b and Fig. 4a). During the preparatory period, the cortical activities converge to a static pattern associated with the upcoming motif, while during motif execution, motif-specific oscillatory activity is generated (Fig. 4b). This recapitulates major experimentally observed features of activity in the motor cortex during movement planning and execution [5, 4, 6].

**Figure 4:**
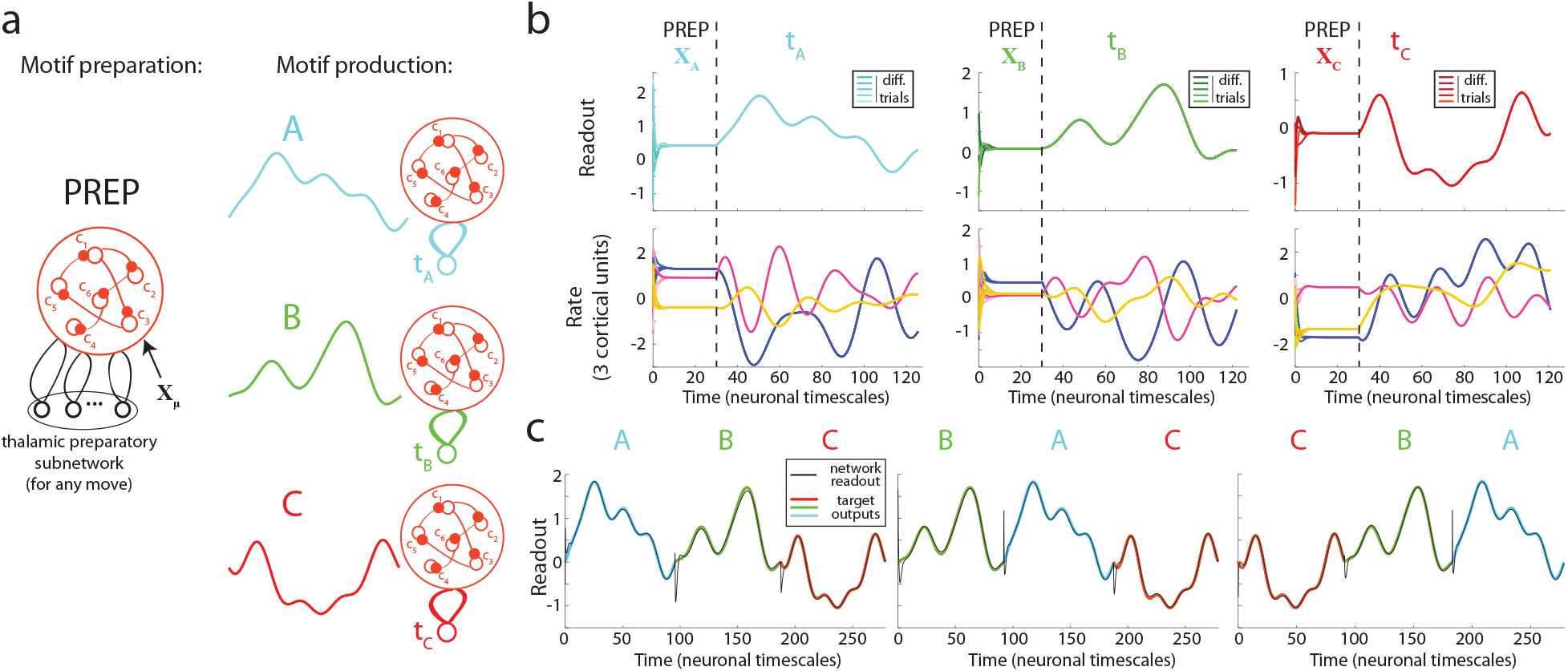
Flexible motor sequencing. **a**) Schematics of motif-specific effective networks and system outputs. The input from the basal ganglia selects thalamic units needed for either motif execution or preparation. During preparation, the cortical population also receives an input specific to the upcoming motif **x**_*μ*_. **b**) Preparation and execution of three example motifs. When the preparatory loop is active, the system output (*upper*) and cortical units (*lower*) quickly converge to the values needed for the upcoming motif. Note that, here, the preparatory period was made much longer than needed to better visualize its effects. Upon activation of the motif-specific loops, the target motifs (*upper*) are composed from rich, oscillatory cortical activities (*lower*). **c**) Generation of sequences. By interleaving brief preparatory periods (here, five cortical timescales) between motif executions, sequences of arbitrary order can begenerated.

Importantly, in our model, the circuit elements corresponding to each motif and to the preparatory period are completely segregated. As a consequence, once a set of motifs is learned, any arbitrary motif sequence order can be generated by the network (Fig. 4c). Furthermore, a new motif can be added to system without interfering with previously learned patterns by setting the connections to and from a new thalamic unit and by adding a new external input during the associated preparatory period.

## 3 Discussion

We have developed a corticothalamocortical model that implements switching linear dynamics. The model can be trained to execute motor motifs by setting its corticothalamic and thalamocortical weights and can switch between these motifs to generate arbitrary sequences. The linearity of the model during motif execution makes it analytically tractable, while the nonlinear inactivation of thalamic units gives it flexibility. Any nonlinear dynamical system can be approximated by switching linear dynamics, with increasing precision (but decreasing tractability) when the switching frequency increases. In the context of motor sequencing, switching is useful to quickly stabilize and adjust the dynamics at each motif transition and to support motif-specific dynamical regimes. In contrast, while linear dynamical systems can have fixed points and interesting transient dynamics [22, 21], they cannot modify their dynamics to suit different motifs. We considered a minimal case of a single thalamic switching unit resulting in a rank-one perturbation through a single corticothalamocortical loop. This allowed us to fully characterize the dynamics with closed-form equations. Importantly, the rank-one perturbation can control multiple eigenvalues. As a result, every configuration of active and inactive thalamic units constructs a thalamocortical network with different dynamics. Thus, the switchable thalamocortical network is equivalent to a large number of different linear networks (Supplementary Fig. S.5). Segregating the units that control the cortical network into their own non-recurrent brain region – the thalamus – simplifies the problems of inputting switch commands into the circuit and of adjusting motif-specific synapses in an efficient and extendable fashion.

Flexible motor sequencing remains a challenge in machine learning [34, 31], because it requires both that new motifs can be learned without destroying previously learned ones [20, 43] and that motif sequences can be generated in orders never experienced during training [2, 14, 34, 31, 37]. Our thalamocortical network overcomes both of these challenges. First, different motifs can be learned with completely separate sets of parameters, preventing interference while still benefiting from the rich dynamics of the shared cortical network. Second, our preparatory network, which is motif independent, ensures that the cortical activity converges to the initial state needed for the upcoming motif, and thus can implement any transition, including novel ones. We show that this convergence can be fast even if the size of the preparatory thalamic population is only a fraction of the size of cortex (Fig. 3). Furthermore, remaining errors in the initial state after preparation have a negligible effect if corticothalamocortical loop weights are optimized for noise robustness. In contrast, artificial neural networks trained with state-of-the-art algorithms suffer both from interference and lack of generalization to new sequences, and can fail catastrophically when presented with these challenges [31, 34, 37].

Our model respects major features of thalamocortical connectivity [49, 26]. However, certain aspects of our model do not match the biology. We assumed (*i*) that, during specific motifs, all cortical units and all disinhibited thalamic units are in a purely linear regime and (*ii*) that thalamus is instantaneous with respect to cortex. In the supplemental materials, we relax these two simplifications by using standard rectified units in place of linear units in both cortex and thalamus and by setting the thalamic time-constant to be ten times faster than cortex rather than infinitely fast. We have verified that the model can still generate motif sequences with dynamics that closely follow our idealized theoretical framework (Supplementary Fig. S.6).

Transitions between motifs in our model are triggered externally by altering the pattern of basal ganglion input to the thalamus. It is likely that the striatum plays this role by inhibiting neurons in the GPi/SNr (the output nuclei of the basal ganglia) at times determined by its overall monitoring of cortical activity. Through its extensive cortical inputs, striatum is well situated for establishing both the timing and selection of motif transitions. In particular, frontal brain regions, which project back to the basal ganglia [33] and whose firing rates can reflect the abstract sequential structure of a task [51, 7, 41], are good candidates for planning and controlling sequence generation in this way.

Our work does not address the mechanisms by which a biologically plausible learning rule could allow the brain to learn the synaptic weights of corticothalamocortical loops. This is somewhat challenging in the model, because the weights must both set the values of eigenvalues and optimize for robustness to noise. The most straightforward interpretation of our model would suggest that plasticity occurs at the level of the synapses of the direct thalamocortical and corticothalamic projections – for which there is some evidence [40, 38, 53, 23, 3]. However, another possibility is that plasticity is situated anywhere within an effective feedforward subnetwork between the thalamic and cortical populations involved in the dynamic production of motor commands. Fine tuning of the synaptic weights between units in the model would be somewhat mitigated by mapping each unit to a population of neurons in a biological network and matching the effective weights between populations [35]. Further work will be needed to investigate whether, in the context of our model, the level of synaptic fine tuning required for generating a complex motor sequence with a realistic number of units can be within a biologically plausible range.

An interesting aspect of our model is the obligatory preparatory period between motifs. A similar preparatory activity appears to be required in motor cortical areas even for the most rapid voluntary movements [5, 29]. Furthermore, as in the data, preparatory activity is approximately orthogonal to the activity in motor production periods [11], a feature also shared by an alternative thalamic preparatory circuit that was recently developed [32].

Our model makes several experimental predictions that can be tested in animals engaged in motor sequence tasks. First, given our proposal that cortical dynamics are controlled via loops through the thalamus which are activated and inactivated by input from the GPi/SNr, we predict that changes in the activity patterns either in thalamus or in the basal ganglia should immediately precede, and be causally related to, changes in cortical dynamics. Recently developed data analysis tools [30] permit the automatic inference of switch times between different dynamical regimes in neural population recordings. Thus simultaneous recordings in motor cortex and either thalamus or GPi/SNr would generate the data needed to test this prediction. Similarly, we predict that switch times in thalamic activity patterns, as well as switch times in cortical dynamics, would reflect points of change in muscle activity and behavior. Next, we predict that perturbative experiments in GPi/SNr and thalamus would have differential effects. A controlled alteration of activity patterns in the basal ganglia could modify sequence structure (i.e., the order of motifs) while leaving individual sequence elements unchanged. Perturbing thalamus, on the other hand, would affect the loops that determine the cortical dynamics and would thus disrupt the execution of individual motifs. Finally, our model also postulates that the thalamic neurons involved in setting cortical dynamics are segregated into motif-specific subpopulations. This prediction could be tested by correlating recordings in thalamus with behavior during a sequence learning task. Interestingly, along these lines, a recent study showed that, when training an animal on two distinct motor tasks, learning for each task is associated with its own synaptic subpopulation in a thalamic-recipient layer of motor cortex [19, 26]. Our interpretation would be that these subpopulations are receiving inputs from motif-specific neurons in thalamus.

In conclusion, our corticothalamocortical model suggests a mechanism for flexible and robust sequence generation, and leads to experimental predictions that can further our understanding of motor system function. In addition, our model reveals how complex cognitive processes such as motor sequencing may rely on neural systems operating on very different timescales. First, slow learning through the adjustment of synaptic weights (e.g., by optimizing some cost function) can construct a library of cognitive building blocks such as motifs. Then, assuming the network architecture is appropriately constrained – as in the case of motif-specific units in the thalamus – the flexible combination and organization of these cognitive building blocks can be achieved online through a selection process like the one that we propose is implemented in the basal ganglia. Understanding the principles of selecting and executing cognitive motifs will be necessary to understand brain function and to build artificial intelligence capable of robust and versatile behavior.

## 4 Methods

Below we present the methods and mathematical results that are needed to build the model. More detailed derivations, and mathematical results that are of interpretative nature, are presented in the Supplementary Material 5.1.

### 4.1 Eigenvalue constraint

Each eigenvalue 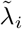 for *i* = 1, 2, …, *N* of 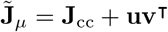 obeys the characteristic equation

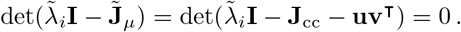

Then, using the matrix determinant lemma under the assumption that 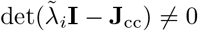,

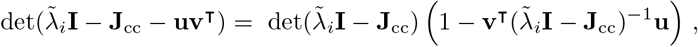

which implies Eq. 6.

If we apply the eigendecomposition of **J**_cc_, we get 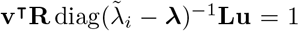. From this we can write the explicit equation for **v** given **u** as

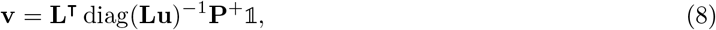

where **P**^+^ is the Moore–Penrose pseudoinverse of **P**.

### 4.2 Definition of the cost function *C* to optimize output noise robustness

We use as our cost function *C* the integrated squared deviation in the network output due to a Gaussian fluctuation ***η*** in the initial conditions with 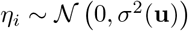:

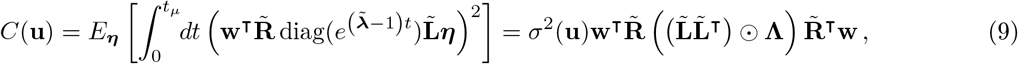

where *E*_***η***_ is an expectation value, *t_μ_* is the duration of the motif, ⊙ is the component-wise Hadamard product, and we have defined the matrix **Λ** with components

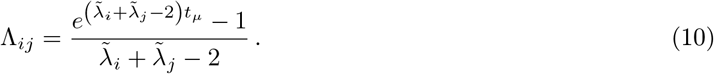

We set the variance of the noise *σ*^2^(**u**) according to the time-averaged squared norm of the activities **c**(*t*) in the absence of noise:

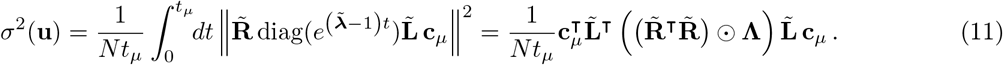

Note that *C*(**u**) and *σ*^2^(**u**) depend on **u** through the eigenvector matrices 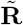 and 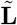.

### 4.3 Efficient computation of the cost function *C*

The cost function *C* given by Eq. 9 requires the eigenvalues 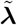 and eigenvectors 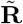 and 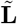 of 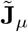. Here we show how these canbecomputed efficiently avoiding explicit eigendecomposition on each iteration of optimization.

First, because 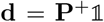, **d** is fully determined by **P** which depends on the eigenvalues of **J**_cc_ and the target eigenvalues, but not on **u**. Given **d**, Eq. 6 can be easily rewritten as a degree-*N* polynomial for 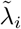 with fixed coefficients, implying that all *N* eigenvalues of 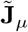 are also fully determined and independent of **u** (Supplementary Sec. 5.1.1). Thus **Λ** can be computed once prior to optimization of Eq. 9.

Next, let 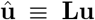 and 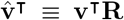. (Note that 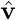 can be calculated from 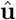 as 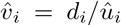.) Then left-multiplying the eigenequation for 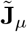 by **L** gives

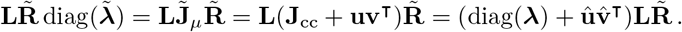

The term corresponding to the *i*^th^ left eigenvector of **J**_cc_ and the *i*^th^ right eigenvector of 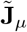 is: 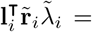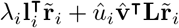. If we assume 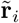 is normalized such that 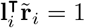 for all *i*, then 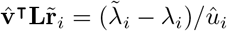 or in matrix form 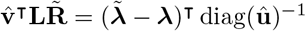. We can substitute this back into the eigenequation to get

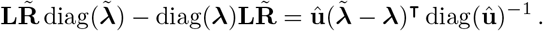

Note that the elements of the left hand side are given as 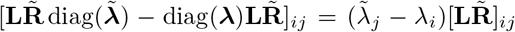. Thus if we define **A** such that 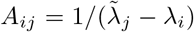, then 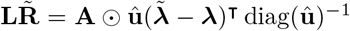. We can rearrange this, drop arbitrary scale factors, and solve for 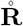 (the unnormalized right eigenvectors of 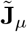) as

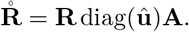

Through a symmetric calculation, the unnormalized left eigenvectors are given as

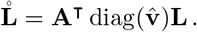

Finally, we normalize the eigenvectors to guarantee that 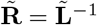 with

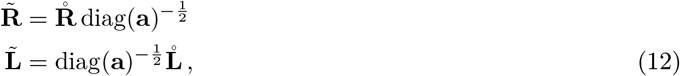

where 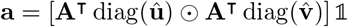.

The eigenvector solutions given in Eq. 12 depend on **R**, **L**, and **A**, all of which can be computed prior to optimization, and on 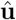 and 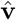 which can be computed from **u** (and **L** and **d**) on each iteration of optimization. Implementing the formulae given in Eq. 12 requires one matrix multiplication each per iteration (in addition to other much cheaper operations). With 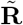, 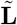, and **Λ** in hand, the remaining operations to compute *C* are dominated by two additional matrix multiplications (in Eqs. 9 & 11). Without the need for more computationally intensive operations (e.g., eigendecomposition or matrix inversion), the cost function *C* is efficient to optimize.

### 4.4 Designing the preparatory network 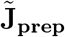

The cost function for determining the matrix 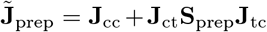 is determined by a calculation similar to the one presented in section 4.2 (and we use the same notation). We want the deviation *δ***c** to go to zero quickly. Its total magnitude over an arbitrarily long preparatory period is

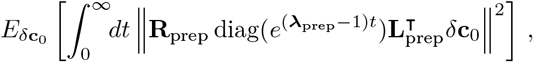

which, if we drop the scale factor given by the variance of *δ***c**_0_, gives the cost function as

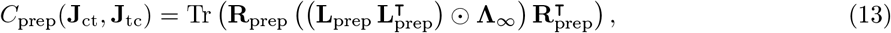

where **Λ**_∞_ is given by Eq. 10 with *t_μ_* → ∞ because the network reaches a fixed point.

After (unconstrained) optimization, we scale the weights according to

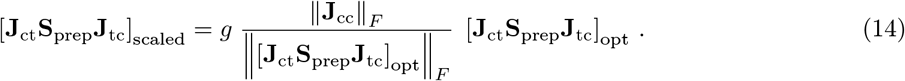

In Fig. 3 we use *g* = 5 which gives weights with reasonable magnitudes and has negligible impact on the decay times of *δ***c** (Fig. 3f) compared to the optimized solution. Indeed, in addition to setting *K* eigenvalues to have large negative real parts (and which do scale with *g*), we find that this corticothalamocortical perturbation also shifts the remaining *N* − *K* eigenvalues away from the Re *λ* = 1 line in a way that is relatively independent of *g* (Fig. 3d). This eliminates any slow mode of the dynamics and guarantees rapid decay of *δ***c** to zero.

### 4.5 Numerical procedures

The simulations were performed using MATLAB and Python. We numerically optimized Eqs. 9 and 4.4 (specifically, the square roots of these cost functions). As discussed in Supplementary Sec. 5.1.2, these equations have multiple local minima. Thus we selected the best among the solutions we found to present as the results in Figs. 2 and 3.

When matchingthedesired output for onemotif *y* to its approximation with asmallnumber of eigenmodes 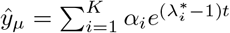, we tried a few different numbers of non-zero eigenmodes *K* (typically between four and twenty, see Fig. S.2). For each value of *K*, we used MATLAB’s fmincon function with 50 different random starts to optimize the real and imaginary part of each amplitude *α_i_* and each eigenvalue 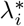 to minimize the mean square difference between *y* and 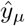. A constraint was added to discourage very large amplitudes or very large real or imaginary part of eigenvalues.

Otherwise stated, we used cortical matrices **J**_cc_ of size *N* = 500, and we made sure that **J**_cc_ − **I** was stable. More specifically, we discarded the matrix if **J**_cc_ − **I** had positive eigenvalues, and we added a small positive value to the leak during numerical simulations of the dynamics to ensure that no instability would arise due to numerical approximations during eigenvalue control with target eigenvalues very close to, or on, the real line.

## Acknowledgements

We thank Christopher J. Cueva, Francesca Mastrogiuseppe and Omri Barak for useful discussions. This research was supported by NIH BRAIN award (U19 NS104649), NSF/NIH Collaborate Research in Computational Neuroscience award (R01 NS105349), NIH Director’s Early Independence award (DP5 OD019897), and NSF NeuroNex award (DBI-1707398), as well as the Leon Levy foundation, the Gatsby Charitable Foundation, and the Swartz Foundation.

