## Supplementary materials for "A model of flexible motor sequencing through thalamic control of cortical dynamics"

### 5 Supplementary Material

#### 5.1 Supplementary Methods

In this article, the network rates obey

$$\dot{\mathbf{c}} = -\mathbf{c} + \left(\mathbf{J}_{\text{cc}} + \mathbf{J}_{\text{ct}}\mathbf{S}_{\mu}\mathbf{J}_{\text{tc}}\right)\mathbf{c} \tag{3}$$

Though similar to the linear version of standard input-driven rate dynamics [50] (i.e., $\dot{\mathbf{c}} = -\mathbf{c} + \mathbf{J}_{\text{cc}}\mathbf{c} + \mathbf{x}_{\mu}$) the motif-specific effect in Eq. 3 appears as an effective change in the connections between units rather than as an additive external input. In other words, our dynamics include a switching nonlinearity at each motif transition, implementing multilinear dynamics whose structure tightly follows the motif sequence.

In this supplementary methods section, we will first lay out derivations and analytical results for this simplified dynamical system in the context of implementing flexible motor sequencing. Then, we will explain how the simplified system can be related to more biologically plausible models of neuronal dynamics. Finally, we will show how major biological constraints can be added to the model without losing the analytical insights from the simplified dynamics and the functionality of the model.

##### 5.1.1 Cortical eigenvalues control through a low-dimensional thalamic modulation

To uncover the full potential of a thalamic modulation of cortical dynamics, we consider the minimal case where a single thalamic unit is left free to interact with cortex during a motif (i.e., there is a single non-zero value in the diagonal matrix $\mathbf{S}_{\mu}$ in Eq. 3). Thus, we can write

$$\dot{\mathbf{c}} = \left(-\mathbf{I} + \mathbf{J}_{\text{cc}} + \mathbf{u}\mathbf{v}^{\intercal}\right)\mathbf{c}\,.$$

We can then ask how much the rank-one perturbation $\mathbf{u}\mathbf{v}^{\intercal}$ can impact the properties of the effective within-motif cortical dynamics, defined by the effective connectivity matrix $\tilde{\mathbf{J}}_{\mu} = \mathbf{J}_{\text{cc}} + \mathbf{u}\mathbf{v}^{\intercal}$. Note that we can set aside the effect of adding the leak $-\mathbf{I}$ to the connectivity, as it simply shifts the eigenvalues by $-1$, without affecting the eigenvectors.

**Control of the eigenvalues of $\tilde{\mathbf{J}}_{\mu}$ through tuning one vector of the rank-one connectivity perturbation** We can write the characteristic polynomial of the effective connectivity matrix $\tilde{\mathbf{J}}_{\mu}$ as a function of properties of $\mathbf{J}_{\text{cc}}$ and of the parameters $\mathbf{u}$ and $\mathbf{v}$. More specifically, we consider the case of an invertible matrix $\mathbf{J}_{\text{cc}}$ whose eigenvalues $\lambda_1 \dots \lambda_j \dots \lambda_N$ are all distinct from one another and from the eigenvalues $\tilde{\lambda}_1 \dots \tilde{\lambda}_i \dots \tilde{\lambda}_N$ of the effective matrix $\tilde{\mathbf{J}}_{\mu}$. Then, the matrix $\mathbf{J}_{\text{cc}} - \tilde{\lambda}_j\mathbf{I}$ is also invertible, which allows us to use the the matrix determinant lemma in order to rewrite the characteristic polynomial for $\tilde{\mathbf{J}}_{\mu}$:

$$0 = \det\left(\mathbf{J}_{\text{cc}} - \tilde{\lambda}_i\,\mathbf{I} + \mathbf{u}\mathbf{v}^{\intercal}\right) = \left(1 + \mathbf{v}^{\intercal}\left(\mathbf{J}_{\text{cc}} - \tilde{\lambda}_i\,\mathbf{I}\right)^{-1}\mathbf{u}\right)\,\det\left(\mathbf{J}_{\text{cc}} - \tilde{\lambda}_i\,\mathbf{I}\right)\,,$$

or

$$1 = \mathbf{v}^{\intercal}\left(\tilde{\lambda}_i\,\mathbf{I} - \mathbf{J}_{\text{cc}}\right)^{-1}\mathbf{u}\,, \tag{6}$$

where in the last step we again used the fact that $\tilde{\lambda}_i$ is not an eigenvalue of $\mathbf{J}_{\text{cc}}$, which implies that $\det\left(\mathbf{J}_{\text{cc}} - \tilde{\lambda}_i\,\mathbf{I}\right) \neq 0$.

We then use the eigendecomposition procedure to write $\mathbf{J}_{\text{cc}} = \mathbf{R}\,\text{diag}(\boldsymbol{\lambda})\mathbf{L}$, where we have concatenated the eigenvalues $\lambda_1, \dots, \lambda_N$ in the vector $\boldsymbol{\lambda}$, and $\mathbf{L}$ and $\mathbf{R}$ are matrices regrouping the left and right eigenvectors of the initial matrix $\mathbf{J}_{\text{cc}}$. This allows us to expand $\left(\tilde{\lambda}_i\,\mathbf{I} - \mathbf{J}_{\text{cc}}\right)^{-1} = \mathbf{R}\,\text{diag}(\tilde{\lambda}_i - \boldsymbol{\lambda})^{-1}\,\mathbf{L}$. We note that $\text{diag}(\tilde{\lambda}_i - \boldsymbol{\lambda})^{-1}$ is a diagonal matrix whose element on the $j^{th}$ row and column is $1/(\tilde{\lambda}_i - \lambda_j)$. Therefore, we can rewrite Eq. 6 as

$$1 = \mathbf{v}^{\intercal}\mathbf{R}\,\text{diag}(\tilde{\lambda}_i - \boldsymbol{\lambda})^{-1}\,\mathbf{L}\mathbf{u}\,. \tag{S.1}$$

Equation S.1 is true for all eigenvalues $\tilde{\lambda}_i$ of $\tilde{\mathbf{J}}_\mu$. As a consequence, in order for the eigenspectrum of $\tilde{\mathbf{J}}_\mu$ to include an ensemble of $K \leq N$ target eigenvalues $\lambda_i^*$, then – with the $K \times N$ matrix $\mathbf{P}$ defined to have elements $P_{ij} = 1/(\lambda_i^* - \lambda_j)$ – the following system of equations holds: $\mathbb{1} = \mathbf{P}\ \mathrm{diag}\left(\mathbf{Lu}\right)\ \mathbf{R}^\intercal\mathbf{v}$. We can therefore write one possible solution for $\mathbf{v}$ as:

$$\mathbf{v} = \mathbf{L}^\intercal\ \mathrm{diag}\left(\mathbf{Lu}\right)^{-1}\ \mathbf{P}^+\ \mathbb{1}\,, \tag{8}$$

where $\mathbf{P}^+$ is the Moore–Penrose pseudoinverse of $\mathbf{P}$ and $\mathbb{1}$ is a vector of all ones. Note that we made the arbitrary choice of setting $\mathbf{v}$ as the determined weight vector in Eq. 8 (while $\mathbf{u}$ can be chosen freely), but we could have equivalently rewritten this equation as a tuning of the vector $\mathbf{u}$.

We can now specify the properties of the fixed cortical matrix $\mathbf{J}_{\mathrm{cc}}$ that are necessary to control eigenvalues according to Eq. 8. First, this equation requires 'inverting' the matrix $\mathbf{P}$, which will lead to accurate solutions if $\mathbf{P}$ is well-conditioned, which will be true if all eigenvalues of $\mathbf{J}_{\mathrm{cc}}$ are distinct enough from each other – as is the case when $\mathbf{J}_{\mathrm{cc}}$ is random. Second, to write Eq. 8, we assumed that $\mathrm{diag}\left(\mathbf{Lu}\right)$ is invertible which just requires $\mathbf{u}$ to not be zero nor orthogonal to one of the left eigenvectors of $\mathbf{J}_{\mathrm{cc}}$ – which is also almost always guaranteed if $\mathbf{u}$ and $\mathbf{J}_{\mathrm{cc}}$ are random.

Hence, Eq. 8 demonstrates that, under general conditions on the cortical matrix $\mathbf{J}_{\mathrm{cc}}$, we can easily find appropriate values for a corticothalamic weight vector $\mathbf{v}$ that will guarantee that the effective matrix $\tilde{\mathbf{J}}_\mu$ contains an ensemble of $K$ desired eigenvalues $\{\tilde{\lambda}_i = \lambda_i^*\}_{i\in[1:K]}$, for $K \leq N$. In practice though, the larger the number of eigenvalues $K$ that we want to control and the further away these desired eigenvalues $\lambda_i^*$ are from the initial spectrum $\{\lambda_j\}_{j\in[1:N]}$, the more fine-tuned the synaptic weights need to be. In extreme cases, the numerical precision becomes insufficient to successfully control the eigenvalues (i.e., the matrix $\mathbf{P}$ becomes ill-conditioned; see Fig. S.1). This can be intuited from the above equation. Indeed, consider the case when $K = N$: computing $\mathbf{v}$ requires taking the classical matrix inverse of $\mathbf{P}$ rather than the pseudoinverse. In addition, if several desired eigenvalues $\lambda_i^*$ are way outside of the ensemble of initial eigenvalues $\lambda_j$, the statistics of the difference $\lambda_i^* - \lambda_j$ will be on a larger scale compared to the differences among the initial eigenvalues, and therefore the matrix $\mathbf{P}$ will have correlated rows and will be hard to invert. Indeed, more formally, in this case, we can write $\lambda_i^* - \lambda_j = \lambda_i^* - (\bar{\lambda} + \epsilon_j)$, with $\epsilon_j << \lambda_i^* - \bar{\lambda}$ and $\bar{\lambda} = \sum_j \lambda_j/N$. Hence, to zeroth order in $\epsilon_j$, $P_{ij} \approx 1/(\tilde{\lambda}_i - \bar{\lambda})$ which is independent of the column index $j$, indicating that different rows will be almost constant and therefore correlated to one another. However, Eq. 8 works very well to force a handful of eigenvalues of the effective matrix $\tilde{\mathbf{J}}_\mu$ to take relevant values corresponding to slow oscillating modes (Fig. S.1).

**Consequences of using $\mathbf{P}^+$ for eigenvalue control on the properties of $\tilde{\mathbf{J}}_\mu$ and of the loop weights**
Note that for any choice of $\mathbf{u}$, if we set $\mathbf{v}$ according to Eq. 8, all $N$ eigenvalues $\tilde{\lambda}_i$ of $\tilde{\mathbf{J}}_\mu$ are fully constrained despite that fact that we only choose $K$ of them (i.e., the $\lambda_i^*$). We can see this because the vector $\mathbf{d}$ in Eq. 8 – defined as $\mathbf{d} \equiv \mathbf{P}^+\mathbb{1} = \mathbf{R}^\intercal\mathbf{v} \odot \mathbf{Lu}$ where $\odot$ is the element-wise product – is independent of $\mathbf{u}$ and because S.1 is true for all $\tilde{\lambda}_i$. Indeed, from S.1 we can write

$$1 = \sum_{j=1}^{N} \frac{d_j}{\tilde{\lambda}_i - \lambda_j} = \frac{\sum_{j=1}^{N} d_j \prod_{k\neq j}\left(\tilde{\lambda}_i - \lambda_k\right)}{\prod_{k=1}^{N}\left(\tilde{\lambda}_i - \lambda_k\right)}\,,$$

which implies

$$0 = \sum_{j=1}^{N}\left(d_j \prod_{k\neq j}\left(\tilde{\lambda}_i - \lambda_k\right)\right) - \prod_{k=1}^{N}\left(\tilde{\lambda}_i - \lambda_k\right)\,. \tag{S.2}$$

Equation S.2 defines the roots of a polynomial where $\tilde{\lambda}_i$ is the variable, and all coefficients are fixed for a given initial matrix $\mathbf{J}_{\mathrm{cc}}$ – which constrains the initial eigenvalues $\lambda_1, \ldots, \lambda_N$ – and for fixed $d_1, \ldots, d_N$ as imposed by Eq. 8. Hence, equation S.2 imposes exactly the $N$ values that $\tilde{\lambda}_i$ can take. In other words, when fixing $\mathbf{d}$ by using the pseudoinverse of $\mathbf{P}$, we constrain the whole eigenspectrum of $\tilde{\mathbf{J}}_\mu$. Moreover, we choose a very particular solution for $\mathbf{d}$: the Moore–Penrose pseudoinverse or the minimal norm solution, which – as can be seen from Eq. S.2 – in turn tends to minimize the maximum magnitude of the coefficients of the characteristic polynomial (except for the leading coefficient associated with $\tilde{\lambda}_i^N$ which is fixed to -1). This further tends to minimize the Lagrange and Cauchy upper bounds on all of the roots of the characteristic polynomial [67]. As a consequence, the modulus of the non-controlled eigenvalues tends to be minimized (Fig. 1d), which is favorable for the stability of the dynamics.

Note that we used the minimum norm, least square error solution for the vector $\mathbf{d}$. Though this indirectly tends to moderate the norm of $\mathbf{u}\mathbf{v}^\intercal$, it is however a slightly different solution from the minimal norm solution for $\mathbf{v}$ (as in [81]). For a given random $\mathbf{u}$, this latter method tends to result in a smaller norm for $\mathbf{v}$. The $N-K$ uncontrolled eigenvalues, however, will tend to have larger modulus than with our approach.

Finally, we remark that when we constrain $\mathbf{d}$, we leave some freedom on the individual norms of the thalamocortical and corticothalamic weight vectors. Indeed, if the vectors $\{\mathbf{u}, \mathbf{v}\}$ solve Eq. 8, then we can define a scale factor $s$ and get $\{\frac{\mathbf{u}}{s}, s\mathbf{v}^\intercal\}$ as another solution. Both of these vector pairs lead to the same value of $\mathbf{d}$ and the same rank-one perturbation $\mathbf{u}\mathbf{v}^\intercal$.

Now that we have described how we fix the eigenspectrum of the effective thalamocortical circuit during a motif, we turn to describing how to improve noise-robustness during motif production.

##### 5.1.2 Improvement of the readout's noise robustness through eigenvector control

In addition to fixing $\mathbf{v}$ using Eq. 8 for eigenvalue control, we also optimize $\mathbf{u}$ in order to improve the noise robustness of the readout with respect to imprecision in the rates at the beginning of the motif – which is the major source of variability in motor cortex [5].

**Necessity for an accurate initial activity pattern at motif start** Our goal is to shape the network's output $\hat{y}$ into a desired motif $\hat{y}_\mu$, which we assume can be composed as a weighted sum of the $K$ eigenmodes built into $\tilde{\mathbf{J}}_\mu$ via Eq. 8: $\hat{y}_\mu(t) = \boldsymbol{\alpha}^\intercal e^{(\tilde{\boldsymbol{\lambda}}-1)t}$, where $t$ is the time elapsed since the start of motif $\mu$ and weight vector $\boldsymbol{\alpha}$ has $\alpha_j = 0$ if $j > K$. (Note that $\hat{y}_\mu$ can be an approximation of an arbitrary target motif $y$; see Supplementary Fig. S.2.) From Eq. 1, we can use the solution for $\mathbf{c}$ and expand into the eigenbasis of $\tilde{\mathbf{J}}_\mu$ to get

$$\hat{y}(t) = \mathbf{w}^\intercal \mathbf{c}(t) = \mathbf{w}^\intercal e^{(\tilde{\mathbf{J}}_\mu - \mathbf{I})t}\mathbf{c}(0) = \mathbf{w}^\intercal \tilde{\mathbf{R}}\,\mathrm{diag}(e^{(\tilde{\boldsymbol{\lambda}}-1)t})\,\tilde{\mathbf{L}}\mathbf{c}(0) = \left[\tilde{\mathbf{L}}\mathbf{c}(0)\right]^\intercal \mathrm{diag}(\mathbf{w}^\intercal\tilde{\mathbf{R}})\, e^{(\tilde{\boldsymbol{\lambda}}-1)t},$$

where the final identity is a rearrangement of vectors. Thus it is clear that $\hat{y}$ will equal $\hat{y}_\mu$ if $\boldsymbol{\alpha} = \mathrm{diag}\left(\mathbf{w}^\intercal\tilde{\mathbf{R}}\right)\tilde{\mathbf{L}}\,\mathbf{c}(0)$, or

$$\mathbf{c}(0) = \mathbf{c}_\mu \equiv \tilde{\mathbf{R}}\,\mathrm{diag}\left(\mathbf{w}^\intercal\tilde{\mathbf{R}}\right)^{-1}\boldsymbol{\alpha}\,, \tag{5}$$

where we define $\mathbf{c}_\mu$ as the ideal initial activity pattern which ensures that $\hat{y} = \hat{y}_\mu$.

Hence, during a preparatory process preceding each motif, the initial rates need to be adjusted such that they approach the desired pattern $\mathbf{c}_\mu$ (through a process described in section 2.4). However, this preparatory process cannot be perfect, and motif production therefore needs to be robust to deviations from the ideal initial pattern $\mathbf{c}_\mu$.

**Definition of the cost function $C$ to optimize output robustness** To guarantee output accuracy under a realistic preparatory period process, we want to adjust the thalamocortical weight vector $\mathbf{u}$ to minimize the expectation of the difference between the readout under ideal initial conditions $\mathbf{c}_\mu$ and the readout under noisy initial conditions $\mathbf{c}_\mu + \boldsymbol{\eta}$. The additive noise $\boldsymbol{\eta}$ contains random i.i.d. elements with zero mean and a variance $\sigma^2(\mathbf{u})$ that is given by the mean squared activity of the cortical rates during the execution of motif $\mu$ in the noiseless setting. More specifically, with $t_\mu$ as the duration of motif $\mu$, we set

$$\begin{aligned}
\sigma^2(\mathbf{u}) &= \frac{1}{t_\mu N}\int_0^{t_\mu} \mathbf{c}^\intercal\,\mathbf{c}\;dt \\
&= \frac{1}{t_\mu N}\mathbf{c}_\mu^\intercal\tilde{\mathbf{L}}^\intercal\left[\int_0^{t_\mu} dt\;\mathrm{diag}\left(e^{(\tilde{\boldsymbol{\lambda}}-1)t}\right)\tilde{\mathbf{R}}^\intercal\,\tilde{\mathbf{R}}\,\mathrm{diag}\left(e^{(\tilde{\boldsymbol{\lambda}}-1)t}\right)\right]\tilde{\mathbf{L}}\mathbf{c}_\mu \\
&= \frac{1}{t_\mu N}\mathbf{c}_\mu^\intercal\tilde{\mathbf{L}}^\intercal\left(\left(\tilde{\mathbf{R}}^\intercal\tilde{\mathbf{R}}\right)\odot\boldsymbol{\Lambda}\right)\tilde{\mathbf{L}}\mathbf{c}_\mu\,,
\end{aligned} \tag{11}$$

where we defined the matrix $\boldsymbol{\Lambda}$ as

$$\Lambda_{ij} = \frac{e^{\left(\tilde{\lambda}_i + \tilde{\lambda}_j - 2\right) t_\mu} - 1}{\tilde{\lambda}_i + \tilde{\lambda}_j - 2} \,. \tag{10}$$

With these, we can write an expression for the cost to be minimized, which sums the effects of the noise $\boldsymbol{\eta}$ on the readout:

$$\begin{aligned} C(\mathbf{u}) &= E_{\boldsymbol{\eta}} \left[ \int_0^{t_\mu} dt \, \left( \mathbf{w}^\intercal e^{\left(\tilde{\mathbf{J}}_\mu - \mathbf{I}\right) t} \mathbf{c}_\mu - \mathbf{w}^\intercal e^{\left(\tilde{\mathbf{J}}_\mu - \mathbf{I}\right) t} \left(\mathbf{c}_\mu + \boldsymbol{\eta}\right) \right)^2 \right] \\ &= E_{\boldsymbol{\eta}} \left[ \int_0^{t_\mu} dt \, \left( \mathbf{w}^\intercal \tilde{\mathbf{R}} \operatorname{diag}\left( e^{\left(\tilde{\boldsymbol{\lambda}} - 1\right) t} \right) \tilde{\mathbf{L}} \, \boldsymbol{\eta} \right)^2 \right] \\ &= \int_0^{t_\mu} dt \, \mathbf{w}^\intercal \tilde{\mathbf{R}} \operatorname{diag}\left( e^{\left(\tilde{\boldsymbol{\lambda}} - 1\right) t} \right) \tilde{\mathbf{L}} \, E_{\boldsymbol{\eta}} \left[ \boldsymbol{\eta} \, \boldsymbol{\eta}^\intercal \right] \tilde{\mathbf{L}}^\intercal \operatorname{diag}\left( e^{\left(\tilde{\boldsymbol{\lambda}} - 1\right) t} \right) \tilde{\mathbf{R}}^\intercal \mathbf{w} \\ &= \sigma^2 \left(\mathbf{u}\right) \mathbf{w}^\intercal \tilde{\mathbf{R}} \left[ \int_0^{t_\mu} dt \, \operatorname{diag}\left( e^{\left(\tilde{\boldsymbol{\lambda}} - 1\right) t} \right) \tilde{\mathbf{L}} \, \tilde{\mathbf{L}}^\intercal \operatorname{diag}\left( e^{\left(\tilde{\boldsymbol{\lambda}} - 1\right) t} \right) \right] \tilde{\mathbf{R}}^\intercal \mathbf{w} \\ &= \sigma^2 \left(\mathbf{u}\right) \, \mathbf{w}^\intercal \tilde{\mathbf{R}} \, \left( \left( \tilde{\mathbf{L}} \, \tilde{\mathbf{L}}^\intercal \right) \odot \boldsymbol{\Lambda} \right) \tilde{\mathbf{R}}^\intercal \mathbf{w} \,. \end{aligned} \tag{9}$$

Our aim is to minimize $C(\mathbf{u})$ by adjusting $\mathbf{u}$, while keeping the vector $\mathbf{v}$ fixed using Eq. 8 to control a set of eigenvalues for the perturbed matrix $\tilde{\mathbf{J}}_\mu = \mathbf{J}_{\mathrm{cc}} + \mathbf{u}\mathbf{v}^\intercal$. During this process, as we discussed previously (Eq. S.2 in Supplementary Sec. 5.1.1), the eigenspectrum of $\tilde{\mathbf{J}}_\mu$ stays fixed, but the eigenvectors of $\tilde{\mathbf{J}}_\mu$ are modified to improve noise robustness.

**Analytic characterization of the cost function** $C$ Equation 9 expresses the cost function $C$ as a function of the properties of the controlled matrix $\tilde{\mathbf{J}}_\mu$: its eigenvectors $\tilde{\mathbf{L}}$ and $\tilde{\mathbf{R}}$, as well as its eigenvalues $\tilde{\lambda}_1, \ldots, \tilde{\lambda}_N$ through the matrix $\boldsymbol{\Lambda}$.

We will now clarify that both $\boldsymbol{\Lambda}$ and these eigenvectors can be expressed as a function of the initial matrix $\mathbf{J}_{\mathrm{cc}}$ and of the perturbation weights $\mathbf{u}$ (and thus $\mathbf{v}$ through Eq. 8). This will reveal interesting properties about nature of the function $C$ and its dependence on the parameters $\mathbf{u}$. Further, these results will reveal that Eq. 9 can be optimized efficiently as there is no need for computing the eigendecomposition of $\tilde{\mathbf{J}}_\mu$ during each step of optimization.

First, as we have noted previously in Supplementary Sec. 5.1.1 the whole eigenspectrum of $\tilde{\mathbf{J}}_\mu$ is fixed because $\mathbf{d}$ and thus the characteristic polynomial for $\tilde{\mathbf{J}}_\mu$ is independent of $\mathbf{u}$ (Eq. S.2). Thus $\boldsymbol{\Lambda}$ can be computed once and for all prior to optimizing Eq. 9.

Second, concerning $\tilde{\mathbf{L}}$ and $\tilde{\mathbf{R}}$, we can use the definition of a left eigenvector $\tilde{\mathbf{l}}_i$ of $\tilde{\mathbf{J}}_\mu = \mathbf{J}_{\mathrm{cc}} + \mathbf{u}\mathbf{v}^\intercal$ to write

$$\tilde{\lambda}_i \, \tilde{\mathbf{l}}_i^\intercal = \tilde{\mathbf{l}}_i^\intercal \left( \mathbf{J}_{\mathrm{cc}} + \mathbf{u}\mathbf{v}^\intercal \right) \qquad \longrightarrow \qquad \tilde{\lambda}_i \, \tilde{\mathbf{l}}_i^\intercal \mathbf{r}_j = \lambda_j \tilde{\mathbf{l}}_i^\intercal \mathbf{r}_j + \tilde{\mathbf{l}}_i^\intercal \mathbf{u}\mathbf{v}^\intercal \mathbf{r}_j \,, \tag{S.3}$$

where we right-multiplied by $\mathbf{r}_j$ in the second step.

We will now consider $\mathring{\mathbf{l}}_i$ which is a particularly scaled version of $\tilde{\mathbf{l}}_i$ satisfying $\mathring{\mathbf{l}}_i^\intercal \, \mathbf{r}_i = 1$. Later, we will take care of renormalizing these arbitrarily scaled eigenvectors $\mathring{\mathbf{l}}_i$ – and $\mathring{\mathbf{r}}_i$ – to get the properly normalized eigenvectors $\tilde{\mathbf{l}}_i^\intercal$ and $\tilde{\mathbf{r}}_i$ such that $\forall i$, $\tilde{\mathbf{l}}_i^\intercal \, \tilde{\mathbf{r}}_i = 1$ as needed. With this scaling, we can rewrite Eq. S.3 for the case of $i = j$ to get

$$\tilde{\lambda}_i = \lambda_i + \mathring{\mathbf{l}}_i^\intercal \mathbf{u} \, \mathbf{v}^\intercal \mathbf{r}_i \qquad \longrightarrow \qquad \mathring{\mathbf{l}}_i^\intercal \mathbf{u} = \frac{\tilde{\lambda}_i - \lambda_i}{\mathbf{v}^\intercal \mathbf{r}_i} \,.$$

Then, substituting this back into Eq. S.3 for $i \neq j$ gives

$$\tilde{\lambda}_i \mathring{\mathbf{l}}_i^\intercal \mathbf{r}_j = \lambda_j \mathring{\mathbf{l}}_i^\intercal \mathbf{r}_j + \frac{\tilde{\lambda}_i - \lambda_i}{\mathbf{v}^\intercal \mathbf{r}_i} \mathbf{v}^\intercal \mathbf{r}_j \qquad \longrightarrow \qquad \mathring{\mathbf{l}}_i^\intercal \mathbf{r}_j = \frac{\mathbf{v}^\intercal \mathbf{r}_j \left( \tilde{\lambda}_i - \lambda_i \right)}{\mathbf{v}^\intercal \mathbf{r}_i \left( \tilde{\lambda}_i - \lambda_j \right)} \,.$$

This relationship is valid for all $j$, so we can use matrix notation to write

$$\mathring{\mathbf{l}}_i^\mathsf{T}\mathbf{R} = \frac{\tilde{\lambda}_i - \lambda_i}{\mathbf{v}^\mathsf{T}\mathbf{r}_i}\,\mathbf{v}^\mathsf{T}\mathbf{R}\,\mathrm{diag}\left(\tilde{\lambda}_i - \boldsymbol{\lambda}\right)^{-1},$$

or

$$\mathring{\mathbf{l}}_i^\mathsf{T} = \frac{\tilde{\lambda}_i - \lambda_i}{\mathbf{v}^\mathsf{T}\mathbf{r}_i}\,\mathbf{v}^\mathsf{T}\mathbf{R}\,\mathrm{diag}\left(\tilde{\lambda}_i - \boldsymbol{\lambda}\right)^{-1}\mathbf{L}\,.$$

Following similar steps, we can also write an equation for the right eigenvectors $\mathring{\mathbf{r}}_i$ that are normalized such that $\mathbf{l}_i^\mathsf{T}\mathring{\mathbf{r}}_i = 1$:

$$\mathring{\mathbf{r}}_i = \frac{\tilde{\lambda}_i - \lambda_i}{\mathbf{l}_i^\mathsf{T}\mathbf{u}}\,\mathbf{R}\,\mathrm{diag}\left(\tilde{\lambda}_i - \boldsymbol{\lambda}\right)^{-1}\mathbf{L}\mathbf{u}.$$

Finally we need to renormalize the eigenvectors such that, $\forall i$, $\tilde{\mathbf{l}}_i^\mathsf{T}\tilde{\mathbf{r}}_i = 1$. We remark that

$$\mathring{\mathbf{l}}_i^\mathsf{T}\mathring{\mathbf{r}}_i = \frac{\left(\tilde{\lambda}_i - \lambda_i\right)^2}{\mathbf{v}^\mathsf{T}\mathbf{r}_i\,\mathbf{l}_i^\mathsf{T}\mathbf{u}}\,\mathbf{v}^\mathsf{T}\mathbf{R}\,\mathrm{diag}\left(\tilde{\lambda}_i - \boldsymbol{\lambda}\right)^{-2}\mathbf{L}\mathbf{u}.$$

Thus, if we define the normalization factor $\beta_i \equiv \left(\mathbf{v}^\mathsf{T}\mathbf{R}\,\mathrm{diag}(\tilde{\lambda}_i - \boldsymbol{\lambda})^{-2}\mathbf{L}\mathbf{u}\right)^{-\frac{1}{2}}$, then we can write the normalized eigenvectors as

$$\begin{cases} \tilde{\mathbf{l}}_i^\mathsf{T} = \dfrac{\mathbf{v}^\mathsf{T}\mathbf{r}_i}{\tilde{\lambda}_i - \lambda_i}\,\beta_i\,\mathring{\mathbf{l}}_i^\mathsf{T} = \beta_i\,\mathbf{v}^\mathsf{T}\mathbf{R}\,\mathrm{diag}\left(\tilde{\lambda}_i - \boldsymbol{\lambda}\right)^{-1}\mathbf{L} \\ \tilde{\mathbf{r}}_i = \dfrac{\mathbf{l}_i^\mathsf{T}\mathbf{u}}{\tilde{\lambda}_i - \lambda_i}\,\beta_i\,\mathring{\mathbf{r}}_i = \beta_i\,\mathbf{R}\,\mathrm{diag}\left(\tilde{\lambda}_i - \boldsymbol{\lambda}\right)^{-1}\mathbf{L}\mathbf{u}\,. \end{cases} \tag{S.4}$$

Note that the eigenvector equations given by Eq. S.4 can be written compactly in matrix form for all $i$ (Eq. 12).

A few interesting conclusions can now be made relative to the optimization of $C$, which depends on the eigenvectors of the perturbed matrix $\tilde{\mathbf{J}}_\mu$ that we now explicitly expressed in Eq. S.4 as a function of *(i)* the properties of the initial matrix $\mathbf{J}_{\mathrm{cc}}$ and *(ii)* the perturbation weights $\mathbf{u}$ and $\mathbf{v}^\mathsf{T}$, both directly and indirectly through the eigenspectrum $\{\tilde{\lambda}_i\}_{1\leq i\leq N}$ of $\tilde{\mathbf{J}}_\mu$ defined by Eq. S.2. First, these equations make it clear that the cost function $C$ can be in general expressed as a ratio of polynomials where the variables are the entries of the weight vector $\mathbf{u}$ that we want to optimize. This implies that there are several local minima in weight space for $C$ (see [74] for a similar result in linear feedforward networks), which justifies the use of a non-local optimization method to minimize $C$ (see 4.5). Second, using these equations, one can avoid implementing a numerical eigendecomposition of $\tilde{\mathbf{J}}_\mu$ at each optimization step – this numerical routine being highly computationally expensive.

**Emergence of eigenvector correlations in the rank-one perturbed cortical matrix** Our equations S.4 for the eigenvectors of $\tilde{\mathbf{J}}_\mu$ clarify some important properties of the dynamics of the effective thalamocortical network. We will focus on the correlations between two eigenvectors $\tilde{\mathbf{r}}_i$ and $\tilde{\mathbf{r}}_j$, as assessed by the cosine of the angle between them $\theta_{ij}$, shown in Fig. 2g and defined as $\cos\theta_{ij} = \mathrm{Re}\left(\tilde{\mathbf{r}}_i^\mathsf{T}\overline{\tilde{\mathbf{r}}_j}\right) / \left(|\tilde{\mathbf{r}}_i|\,|\tilde{\mathbf{r}}_j|\right)$. These correlations have indeed been shown to create non-intuitive and complex non-normal amplification dynamics in linear networks [12, 36, 21].

The value $\cos\theta_{ij}$ approaches 1 (or, equivalently, the eigenvectors are almost parallel) if the real and imaginary parts of the two eigenvectors have very similar directions, irrespective of the norm of these vectors. Hence, from Eq. S.4, we can predict that $\cos\theta_{ij}$ will be large for the vectors $\tilde{\mathbf{r}}_i$ and $\tilde{\mathbf{r}}_j$ if the vectors $\mathbf{R}\,\mathrm{diag}\left(\tilde{\lambda}_i - \boldsymbol{\lambda}\right)^{-1}\mathbf{L}\mathbf{u}$ and $\mathbf{R}\,\mathrm{diag}\left(\tilde{\lambda}_j - \boldsymbol{\lambda}\right)^{-1}\mathbf{L}\mathbf{u}$ have similar directions. This happens when $\tilde{\lambda}_i$ and $\tilde{\lambda}_j$ are much closer to one another than to eigenvalues in the initial spectrum. Notably, if controlled eigenvalues are far from the initial spectrum their eigenvector correlations will be high (Figs. 1d and 2g). But this is also seen for uncontrolled eigenvalues that happen to be very close to one another even at the center of the eigenvalue distribution (Supplementary Fig. S.3a,b).

The above-mentioned significant eigenvector correlations are responsible for the non-normal amplification effects in our network (Fig. 2), and are in stark contrast to the correlations we see in other classes of matrices that we study as controls. In Fig. 2h-i we consider two classes of control matrices – each of the form $\mathbf{R}_c \operatorname{diag}\left(\tilde{\boldsymbol{\lambda}}\right)\mathbf{R}_c^{-1}$ – where the eigenvector matrices $\mathbf{R}_c$ were generated either from a random Gaussian matrix or an normal matrix:

- We created random Gaussian matrices from which we selected those with the same number of real eigenvalues as their matched $\tilde{\mathbf{J}}_\mu$. We then extracted their eigenvectors through eigendecomposition, and finally aligned real eigenvectors to real eigenvalues and complex conjugate eigenvector pairs to complex conjugate eigenvalue pairs of $\tilde{\boldsymbol{\lambda}}$.

- We created random orthogonal eigenvectors with appropriate numbers of real and complex conjugate pairs using the method described in [76]. We first created $N$ random real orthogonal eigenvectors $\mathbf{R}_c^{\mathrm{Re}}$ using the QR-decomposition based methodology developped in [76] to uniformly sample the orthogonal group. We then created the appropriate $n_{\mathrm{Im}}$ number of complementary pairs of complex conjugate eigenvectors $\mathbf{R}_c^{\mathrm{Im}}$ through multiplication of an $n_{\mathrm{Im}}$ subset of the columns of $\mathbf{R}_c^{\mathrm{Re}}$, by the eigenvectors of an $n_{\mathrm{Im}}$ dimensional real random orthogonal matrix created with the same methodology and selected to only have complex conjugate eigenvalues (which was very common). Then, by concatenating the $N - n_{\mathrm{Im}}$ columns of $\mathbf{R}_c^{\mathrm{Re}}$ that we had left aside, with the complementary $n_{\mathrm{Im}}$ columns of $\mathbf{R}_c^{\mathrm{Im}}$, we could create a complete set of random orthogonal eigenvectors with the appropriate number of real and complex conjugate columns.

The first type of control matrix will have some limited amount of eigenvector correlations, leading to relatively little non-normal amplification. However, it still leads to slightly slower activity decay – and therefore larger activities – compared to orthogonal eigenvectors (see Fig. 2h-i; see also [56]).

Indeed, with the second type of eigenvector matrix which is fully orthogonal, no non-normal amplification occurs. More specifically, in the case of orthogonal eigenvectors, one can show that the expected integral of the square activity norm $a^{\mathrm{orth}} \equiv E_{\delta\mathbf{c}_0} \int dt\, \|\mathbf{c}\|^2$ only depends on the real part of the eigenvalues, according to

$$a^{\mathrm{orth}} = E_{\delta\mathbf{c}_0}\left[\int_0^{t_\mu} dt\, \mathbf{c}^{\mathsf{T}}\, \mathbf{c}\right] = E_{\delta\mathbf{c}_0}\left[\int_0^{t_\mu} dt\ \operatorname{Tr}\left(\mathbf{c}\,\mathbf{c}^H\right)\right] = \epsilon^2 \sum_i \frac{e^{2\operatorname{Re}\left(\tilde{\lambda}_i - 1\right) t_\mu} - 1}{2\operatorname{Re}\left(\tilde{\lambda}_i - 1\right)}\,, \tag{S.5}$$

where $\epsilon^2$ is the variance of the initial i.i.d. rates $\delta\mathbf{c}_0$ and the superscript $H$ indicates conjugate transposition. Notice that in Eq. S.5, the different timescales of eigenmodes (i.e., $1/\operatorname{Re}(\tilde{\lambda}_i - 1)$) are simply added together and do not otherwise interact. In contrast, non-normal amplification is a deviation of the expected square activity norm from equation S.5 due to eigenvector correlations:

$$a^{\mathrm{corr}} = \epsilon^2 \operatorname{Tr}\left[\tilde{\mathbf{R}}\left(\left(\tilde{\mathbf{L}}\,\tilde{\mathbf{L}}^H\right) \odot \mathbf{K}\right)\tilde{\mathbf{R}}^H\right]\,, \tag{S.6}$$

where we follow a similar derivation as in Eq. 9, and we defined the matrix $\mathbf{K}$ with elements $K_{ij} = (e^{(\tilde{\lambda}_i + \tilde{\lambda}_j^H - 2)t_\mu} - 1)/(\tilde{\lambda}_i + \tilde{\lambda}_j^H - 2)$. Here we use the conjugate transpose $\mathbf{c}^H$ (instead of $\mathbf{c}^{\mathsf{T}}$ as in Eq. 9), and see that the equation does not simplify. Similarly, if we had used the conjugate transpose when deriving Eq. 9, we would have seen no simpler form.

When comparing Eqs. S.5 and S.6, it is clear that the correlated eigenvectors in Eq. S.6 lead to magnified interactions between the dynamics of eigenmodes which combine into the neural activities: pairs of eigenvalues interact in the matrix $\mathbf{K}$, and these interaction terms are multiplied by large numbers contained in the matrices $\tilde{\mathbf{L}}$ and $\tilde{\mathbf{R}}$ arising because these ill-conditioned, correlated matrices are the inverses of one another.

###### 5.1.3 Designing a thalamus subnetwork to prepare cortex for upcoming motif production

To transition between two motifs during a "preparatory period", we need to accomplish two goals. First, we want the rates to converge towards a desired initial state $\mathbf{c}_\mu$ (given by Eq. 5) which will permit the production of the upcoming motif $\mu$, and second we want this convergence to occur as rapidly as possible.

We can assume that the corticothalamic circuit during the preparatory period is given as $\tilde{\mathbf{J}}_{\mathrm{prep}} = \mathbf{J}_{\mathrm{cc}} + \mathbf{J}_{\mathrm{ct}}\mathbf{S}_{\mathrm{prep}}\mathbf{J}_{\mathrm{tc}}$, where $\mathbf{S}_{\mathrm{prep}}$ selects an ensemble of thalamic units that are specific to the preparatory period.

Then we can force the network to converge towards $\mathbf{c}_\mu$ by using a static external cortical input $\mathbf{x}_\mu$ that we tailor for motif $\mu$. Specifically, if the network dynamics are

$$\dot{\mathbf{c}} = \left(\tilde{\mathbf{J}}_{\text{prep}} - \mathbf{I}\right) \mathbf{c} + \mathbf{x}_\mu \,,$$

then, we want to set $\mathbf{x}_\mu$ such that when $\dot{\mathbf{c}} = 0$, $\mathbf{c} = \mathbf{c}_\mu$. This is satisfied if

$$\mathbf{x}_\mu = -\left(\tilde{\mathbf{J}}_{\text{prep}} - \mathbf{I}\right) \mathbf{c}_\mu \,. \tag{S.7}$$

In order to make the preparatory period as fast as possible, we seek to find a corticothalamocortical connectivity $\mathbf{J}_{\text{ct}}\mathbf{S}_{\text{prep}}\mathbf{J}_{\text{tc}}$ for which the difference between the rates and the steady-state $\delta\mathbf{c} = \mathbf{c} - \mathbf{c}_\mu$ goes to zero as fast as possible. The dynamics of $\delta\mathbf{c}$ read as

$$\dot{\delta\mathbf{c}} = \left(\tilde{\mathbf{J}}_{\text{prep}} - \mathbf{I}\right) \delta\mathbf{c} \qquad \longrightarrow \qquad \delta\mathbf{c}(t) = \mathbf{R}_{\text{prep}}\, \text{diag}(e^{(\boldsymbol{\lambda}_{\text{prep}}-1)t})\, \mathbf{L}_{\text{prep}}\, \delta\mathbf{c}_0 \,,$$

where $\delta\mathbf{c}_0$ is the starting state of the preparatory period, $\mathbf{R}_{\text{prep}}$ and $\mathbf{L}_{\text{prep}}$ are the right and left eigenvector matrices of $\tilde{\mathbf{J}}_{\text{prep}}$ respectively, and $\boldsymbol{\lambda}_{\text{prep}}$ is a vector containing the eigenvalues of $\tilde{\mathbf{J}}_{\text{prep}}$. The starting state of the preparatory period $\delta\mathbf{c}_0$ depends on the ending state of the previous motif which we assume is unknown. Therefore, to find a fast corticothalamic connectivity, we construct a cost function that minimizes $\delta\mathbf{c}$ while averaging over $\delta\mathbf{c}_0$. Specifically, we minimize the squared norm of $\delta\mathbf{c}$ and take $\delta\mathbf{c}_0$ to be a vector of uncorrelated random variables with zero mean and standard deviation $\sigma(\delta\mathbf{c}_0)$:

$$\begin{aligned}
&C_{\text{prep}}(\mathbf{J}_{\text{ct}}\mathbf{S}_{\text{prep}}\mathbf{J}_{\text{tc}}) \\
&\qquad = E_{\delta\mathbf{c}_0}\left[\int_0^\infty \|\delta\mathbf{c}\,(t)\|^2\, dt\right] \\
&\qquad = \int_0^\infty dt\ \text{Tr}\left(\mathbf{R}_{\text{prep}}\ \text{diag}\left(e^{(\boldsymbol{\lambda}_{\text{prep}}-1)\,t}\right)\ \mathbf{L}_{\text{prep}}\ E_{\delta\mathbf{c}_0}\left[\delta\mathbf{c}_0\ \delta\mathbf{c}_0^{\intercal}\right]\ \mathbf{L}_{\text{prep}}^{\intercal}\ \text{diag}\left(e^{(\boldsymbol{\lambda}_{\text{prep}}-1)\,t}\right)\ \mathbf{R}_{\text{prep}}^{\intercal}\right) \\
&\qquad = \sigma(\delta\mathbf{c}_0)\ \text{Tr}\left(\mathbf{R}_{\text{prep}}\left(\left(\mathbf{L}_{\text{prep}}\,\mathbf{L}_{\text{prep}}^{\intercal}\right)\odot\mathbf{A}\right)\mathbf{R}_{\text{prep}}^{\intercal}\right),
\end{aligned} \tag{S.8}$$

where we defined the matrix $A_{ij} = 1/(2 - \lambda_{\text{prep},i} - \lambda_{\text{prep},j})$. Note that $\mathbf{A} = \lim_{t_\mu\to\infty}\boldsymbol{\Lambda}$ where $\boldsymbol{\Lambda}$ was defined in Eq. 10.

When optimizing this cost function, we did not use any mechanism to constrain the norm of the synaptic weights. However, we observed that the numerical optimization did not appear to drift towards more and more negative weights, and instead appeared to slow down around a solution with relatively large but finite (i.e., much smaller than machine limit) perturbation weights. This suggests that the optimization was not attempting to converge towards an infinitely large perturbation which would set all effective eigenvalues to $-\infty$; indeed, while theoretically feasible if a very large precision of the weights could be achieved, this type of perturbation matrix would also lead to very large eigenvector correlations which slow down the dynamics and can counteract the effect of very negative eigenvalues [56].

Further hints that the solution we found is not an approximation of an 'infinitely large eigenvalues' solution can be seen from its behavior in response to scaling of the optimized perturbation weights $[\mathbf{J}_{\text{ct}}\mathbf{S}_{\text{prep}}\mathbf{J}_{\text{tc}}]_{\text{opt}}$. We investigated this because these weights, while well beneath the machine limit, are still relatively large, raising the question of biological plausibility. More specifically, we observed that scaling the Frobenius norm of the optimized solution as per

$$[\mathbf{J}_{\text{ct}}\mathbf{S}_{\text{prep}}\mathbf{J}_{\text{tc}}]_{\text{scaled}} = g\ \frac{||\mathbf{J}_{\text{cc}}||_F}{\left|\left|[\mathbf{J}_{\text{ct}}\mathbf{S}_{\text{prep}}\mathbf{J}_{\text{tc}}]_{\text{opt}}\right|\right|_F}\ [\mathbf{J}_{\text{ct}}\mathbf{S}_{\text{prep}}\mathbf{J}_{\text{tc}}]_{\text{opt}}$$

has a negligible impact on the cost $C_{\text{prep}}$ as long as $g$ is not too small. More specifically, varying $g$ from its maximum value of $g = \left\|\,[\mathbf{J}_{\text{ct}}\mathbf{S}_{\text{prep}}\mathbf{J}_{\text{tc}}]_{\text{opt}}\,\right\|_F / \,\|\mathbf{J}_{\text{cc}}\|_F$ to $g = 5$ (as in Fig. 3) virtually does not change the convergence times of the thalamocortical network as shown in Fig. 3f. More specifically, we observed that the eigenspecturm of the preparatory network $\boldsymbol{\lambda}_{\text{prep}}$ is composed of two classes of eigenvalues (see Fig. 3b). First, assuming that the rank of $\mathbf{J}_{\text{ct}}\mathbf{S}_{\text{prep}}\mathbf{J}_{\text{tc}}$ is $M$, $\boldsymbol{\lambda}_{\text{prep}}$ contains $M$ eigenvalues whose values are very close to the values of the $M$ non-zero eigenvalues of $[\mathbf{J}_{\mathrm{ct}}\mathbf{S}_{\mathrm{prep}}\mathbf{J}_{\mathrm{tc}}]_{\mathrm{scaled}}$. For these eigenvalues, the real parts are increasingly negative as $g$ increases. The $N-M$ eigenvalues in the second class are observed to shift away from the line $\mathrm{Re}\,\lambda = 1$ towards more negative values compared to the eigenspectrum of $\mathbf{J}_{\mathrm{cc}}$. Furthermore, this shift appears to be relatively independent of $g$. This second class is the one which is most instrumental in driving fast convergence as it effectively eliminates any slow mode of the dynamics, and its insensitivity over a large range of $g$ – possibly because of the preservation of the structure in the correlation between the values of $\mathbf{J}_{\mathrm{cc}}$ and those of $[\mathbf{J}_{\mathrm{ct}}\mathbf{S}_{\mathrm{prep}}\mathbf{J}_{\mathrm{tc}}]_{\mathrm{scaled}}$ – explains the fast convergence of the cortical activities towards steady-state over these different $g$ values.

###### 5.1.4 Relating switching multilinear dynamics to neuronal population dynamics equations

Here we show that the switching multilinear dynamics that we consider in the main text can be related to more biologically plausible (positive) population rate equations.

Indeed, the firing rate in a population of spiking neurons can be mathematically well-approximated using a simple two-stage process that can be mapped onto firing rate equations [78]. First, the synaptic inputs are linearly filtered – for instance, for Leaky-Integrate-and-Fire neurons, with an exponential filter through feeding the input into a first order linear differential equation – which gives rise to a 'voltage-like' variable. Second, this 'voltage-like' variable is passed through a static non-linearity. This static non-linearity can be qualitatively described as rectified-linear above a certain threshold value $\boldsymbol{\theta}$ of the voltage-like variable $\mathbf{v}$ (symbolized as $[\mathbf{v}-\boldsymbol{\theta}]^{+}$).

Following this framework, here, we consider a cortical network consisting of populations with 'voltage-like' variables $\mathbf{v}$ recurrently interacting through the effective connectivity matrix $\mathbf{J}_{\mathrm{cc}}$, with resting voltage $\mathbf{v_{rest}}$, and responding to an effective input $\mathbf{p}_\mu$ which stays constant during the motif $\mu$:

$$\dot{\mathbf{v}} = -\left(\mathbf{v}-\mathbf{v_{rest}}\right) + \mathbf{J}_{\mathrm{cc}}\,\left[\mathbf{v}-\boldsymbol{\theta}\right]^{+} \;+\; \mathbf{p}_\mu \tag{S.9}$$

Note that the recurrent connectivity matrix $\mathbf{J}_{\mathrm{cc}}$ is 'effective' in the sense that populations can impact one another both positively and negatively. This effective connectivity can be mapped to a biological network with separated inhibition and excitation [65], by assuming that each unit described by equation S.9 actually maps to two subpopulations: one with excitatory neurons which can have a slower effective timescale [83, 72], and another one with just inhibitory neurons that have faster dynamics [75]. Then, effective excitatory connections between populations naturally map to projections from the excitatory neurons of the sending populations to the excitatory neurons of the respective receiving populations. In addition, effective inhibitory connections between populations can be mapped to a disynaptic pathway involving the excitatory neurons of the sending populations projecting towards the inhibitory neurons of the receiving populations, and these inhibitory neurons locally inhibiting their respective excitatory 'neighbors' [65].

We can then define $\mathbf{c_{abs}} = (\mathbf{v}-\boldsymbol{\theta})$, and assume that the dynamics stay in the linear regime as the input and recurrent connections keep $\mathbf{v}$ above threshold, to get:

$$\dot{\mathbf{c}}_{\mathbf{abs}} = -\mathbf{c_{abs}} - \boldsymbol{\theta} + \mathbf{v_{rest}} + \mathbf{J}_{\mathrm{cc}}\,\mathbf{c_{abs}} \;+\; \mathbf{p}_\mu$$

We can also define $\bar{\mathbf{c}} = \left(\mathbf{I}-\mathbf{J}_{\mathrm{cc}}\right)^{-1}\,\left(\mathbf{v_{rest}}-\boldsymbol{\theta}+\mathbf{p}_\mu\right)$ as the steady-state value for $\mathbf{c_{abs}}$ (which can be found by solving $\dot{\mathbf{c}}_{\mathbf{abs}} = 0$), which allows us to write:

$$\dot{\mathbf{c}}_{\mathbf{abs}} = -\left(\mathbf{c_{abs}}-\bar{\mathbf{c}}\right) + \mathbf{J}_{\mathrm{cc}}\,\left(\mathbf{c_{abs}}-\bar{\mathbf{c}}\right)$$

This is the model that we consider for the motor cortical firing rates, with the addition of an input from the thalamus (with rates $[\mathbf{t_{abs}}]^{+}$) through the thalamocortical weights $\mathbf{J}_{\mathrm{ct}}$:

$$\dot{\mathbf{c}}_{\mathbf{abs}} = -\left(\mathbf{c_{abs}}-\bar{\mathbf{c}}\right) \;+\; \mathbf{J}_{\mathrm{cc}}\,\left(\mathbf{c_{abs}}-\bar{\mathbf{c}}\right) \;+\; \mathbf{J}_{\mathrm{ct}}\,\left[\mathbf{t_{abs}}\right]^{+} \tag{S.10}$$

Following a similar model for the population firing rate of spiking neurons in the thalamus as for cortex [78], we define the thalamic rates $[\mathbf{t_{abs}}]^{+} = [\mathbf{v_{thal}}-\boldsymbol{\theta}_{\mathbf{thal}}]^{+}$ as the rectification of an 'activation' voltage-like variable $\mathbf{t}$ itself undergoing linear dynamics which, in absence of input, revolve around a baseline $\bar{\mathbf{t}}$:

$$\tau_t\,\dot{\mathbf{t}}_{\mathbf{abs}} = -\left(\mathbf{t_{abs}}-\bar{\mathbf{t}}\right) + \mathbf{J}_{\mathrm{tc}}\,\mathbf{c_{abs}} + \mathbf{q}_\mu \tag{S.11}$$

The inputs to thalamus include cortical projections through the effective weights $\mathbf{J}_{\text{tc}}$. Positive cortico-thalamic connections can occur through direct projections from cortex, while inhibitory effective connections can correspond to an indirect cortical projection relayed by the reticular thalamic nucleus [54]. In addition, thalamus receives an input from the basal ganglia $\mathbf{q}_\mu$.

We consider the case when the basal ganglia drive to thalamus consists of either sustained strong inhibition during a motif, or a complete release from inhibition (zero drive) in a small subset of thalamic neurons which we will index as rows with the value 1 in the diagonal of an otherwise empty matrix $\mathbf{S}_\mu$. In addition, we assume that these released thalamic neurons then interact with cortex within the linear regime.

Then, we can write:

$$[\mathbf{t_{abs}}]^+ = \mathbf{S}_\mu\, \mathbf{t_{abs}}$$

Taking these assumptions together, we get the following system of equations for the thalamocortical network:

$$\begin{cases} \dot{\mathbf{c}}_{\mathbf{abs}} = -\left(\mathbf{c_{abs}} - \bar{\mathbf{c}}\right) \,+\, \mathbf{J}_{\text{cc}}\, \left(\mathbf{c_{abs}} - \bar{\mathbf{c}}\right) \,+\, \mathbf{J}_{\text{ct}}\, \mathbf{S}_\mu\, \mathbf{t_{abs}} \\ \tau_t\, \mathbf{S}_\mu\, \dot{\mathbf{t}}_{\mathbf{abs}} = -\,\mathbf{S}_\mu\, \left(\mathbf{t_{abs}} - \bar{\mathbf{t}}\right) + \,\mathbf{S}_\mu\, \mathbf{J}_{\text{tc}}\, \mathbf{c_{abs}} \end{cases} \tag{S.12}$$

Finally, in the main article, we consider the limit in which thalamus is very fast relative to cortex ($\tau_t \to 0$). Indeed, thalamus lacks recurrent excitation, making it react at the relatively fast timescales that are intrinsic to a single-neuron, while the recurrently connected cortical populations can be much slower [83, 72]. Hence, we assume that thalamus almost instantly follows its input, implying:

$$\mathbf{S}_\mu\, \mathbf{t_{abs}} = \mathbf{S}_\mu\, \bar{\mathbf{t}} \,+\, \mathbf{S}_\mu\, \mathbf{J}_{\text{tc}}\, \mathbf{c_{abs}} \tag{S.13}$$

To further simplify the equation, we define:

$$\mathbf{c} = \mathbf{c_{abs}} -\, \left(\mathbf{J}_{\text{cc}} + \mathbf{J}_{\text{ct}}\, \mathbf{S}_\mu \mathbf{J}_{\text{tc}} - \mathbf{I}\right)^{-1}\, \left((\mathbf{I} - \mathbf{J}_{\text{cc}})\, \bar{\mathbf{c}} + \mathbf{J}_{\text{ct}}\, \mathbf{S}_\mu\, \bar{\mathbf{t}}\right) := \mathbf{c_{abs}} - \mathbf{a}_\mu \tag{S.14}$$

We can then finally write, from equations S.12, S.13 and S.14:

$$\dot{\mathbf{c}} = -\mathbf{c} + \left(\mathbf{J}_{\text{cc}} + \mathbf{J}_{\text{ct}} \mathbf{S}_\mu \mathbf{J}_{\text{tc}}\right)\, \mathbf{c} \tag{S.15}$$

This is the equation for the cortical rates $\mathbf{c}$ that we used throughout the main text, that we then introduced with the help a centered rectified thalamic variable $\mathbf{t} = \mathbf{S}_\mu\, \mathbf{t_{abs}} - \mathbf{S}_\mu \mathbf{J}_{\text{tc}} \mathbf{c_{abs}}$ in Eq. 2. We stress that this means that the target readout patterns that we are displaying in the main text are deviations from a motif-specific baseline readout value $\mathbf{w}^\intercal\, \mathbf{a}_\mu$.

Note that our framework could be generalized in several ways. First, there could be additional, non-plastic thalamic loops that could interact with cortex – either in a motif-specific fashion or constantly – and that would simply take part, along with cortex, in an effective 'fixed' recurrent network that the plastic thalamic loop would have to modulate during a motif. Second, even though we only considered here that all the 'plastic' thalamic loops that are not involved in producing the current motif are shut off by basal ganglia, the model may be extended to any scenario where the dynamics of the non-plastic neurons are dominated by constant firing during the motif (e.g., by strong inputs from other intra- and extra-thalamic sources that dominate over the inputs from cortex). Indeed, these driven thalamic neurons would then act as an external constant input added to the effective cortical dynamics, whose effect could therefore easily be canceled by simple substraction of an effective copy of this signal on the readout.

Finally, while we have used two additional assumptions compared to the initial population rate equations Eq. S.9 – namely instantaneous thalamus and linearization of the dynamics with the exception of basal-ganglia driven rectification of thalamus – we will actually show that we can relax those assumptions in the following sections.

###### 5.1.5 Introducing major biological constraints: rate positivity and realistic thalamic timescale

In the main text, we use simplified equations for the rate dynamics (Eq. 3) which make two major approximations relative to a more biologically-inspired dynamics [78]: we assumed that *(i)* the thalamic units respond instantaneously to their inputs, and *(ii)* the rates' dynamics could be linearized and centered around baseline values. Here, we show that we can relax those assumptions while still shaping the readout of our network in a sequence of motifs, and while staying very close to the theoretical framework that we describe in the main text.

More specifically, we will still constrain our dynamics such that for a particular motif, our theory (which assumes an instantaneous thalamus) is still used to set the effective eigenspectrum (through the tuning of the corticothalamic weights $\mathbf{v}$ in Eq. 8) and the initial centered rates $\mathbf{c}_\mu$ (as a function of the desired amplitudes $\boldsymbol{\alpha}_\mu$ for the eigenmodes; eq 5):

$$\begin{cases} \mathbf{v} = \mathbf{L}^\intercal \operatorname{diag}(\mathbf{L}\mathbf{u})^{-1} \mathbf{P}^{+} \mathbb{1} & (8) \\ \mathbf{c}(0) = \mathbf{c}_\mu \equiv \tilde{\mathbf{R}} \operatorname{diag}\left(\tilde{\mathbf{R}}^\intercal \mathbf{w}\right)^{-1} \boldsymbol{\alpha}_\mu \,, & (5) \end{cases}$$

where $\mathbf{u}$ is the thalamocortical weight vector from the single thalamic unit interacting with cortex during a motif, and $\mathbf{v}^\intercal$ is the corresponding corticothalamic weight vector.

In the following, however, thalamus will only be assumed to be reasonably fast: ten times faster than cortex. This means that the delay in thalamus will induce deviations that, as we will show, are small enough to be treated as a correlated and biased noise introduced in the ideal instantaneous model. To minimize the effect of this biased noise, we will consider the full dynamical model including thalamus and positive rates. We will start with linearized and centered equations to first explain how to mitigate the effect of the thalamic delay. We will then later treat the issues of rates positivity and rectifying non-linearity.

We start from Eq. S.10 to describe the cortical dynamics, and also to describe the dynamics in the subset of the thalamic units that are in the linear regime and interacting with cortex (indexed by the 'act' index, and corresponding to a subset of units from Eq. S.11; note that we dropped the 'abs' index for conciseness). We can then write:

$$\begin{cases} \dot{\mathbf{c}_{\mathbf{abs}}} = -\left(\mathbf{c}_{\mathbf{abs}} - \bar{\mathbf{c}}\right) + \mathbf{J}_{\mathrm{cc}}\left(\mathbf{c}_{\mathbf{abs}} - \bar{\mathbf{c}}\right) + \mathbf{J}_{\mathrm{ct}}^{\mathrm{act}}\, \mathbf{t}_{\mathbf{act}} \\ \dot{\mathbf{t}_{\mathbf{act}}} = \dfrac{1}{\tau_t}\left[-\left(\mathbf{t}_{\mathbf{act}} - \overline{\mathbf{t}_{\mathbf{act}}}\right) + \mathbf{J}_{\mathrm{tc}}^{\mathrm{act}}\, \mathbf{c}_{\mathbf{abs}}\right] \end{cases} \tag{S.16}$$

We remind the reader that, implicitly, the vectors $\bar{\mathbf{c}}$ and $\overline{\mathbf{t}_{\mathbf{act}}}$ include an external input which is specific to the current motif during motif execution, or to the upcoming motif during motif transition, in line with the framework described in the main text.

We can then express the dynamics using a single vector $\mathbf{z}_{\mathbf{abs}}$ to concatenate both the cortical and non-rectified thalamic rates: for $i \leq N$, $[\mathbf{z}_{\mathbf{abs}}]_i = [\mathbf{c}_{\mathbf{abs}}]_i$, and for $i > N$, $[\mathbf{z}_{\mathbf{abs}}]_i = [\mathbf{t}_{\mathbf{act}}]_i$. Similarly, we define a concatenated vector of biases $\bar{\mathbf{z}}$ regrouping $\bar{\mathbf{c}}$ and $\overline{\mathbf{t}_{\mathbf{act}}}$. Hence, we can write:

$$\dot{\mathbf{z}_{\mathbf{abs}}} = \mathbf{M}_{\mathbf{eff}}\, \mathbf{z}_{\mathbf{abs}} + \mathbf{V}_{\mathbf{eff}}\, \bar{\mathbf{z}}\,, \tag{S.17}$$

where we defined the matrices $\mathbf{M}_{\mathbf{eff}}$ and $\mathbf{V}_{\mathbf{eff}}$ such that:

- for $i \leq N$ and $j \leq N$, $[\mathbf{M}_{\mathbf{eff}}]_{i,j} = [\mathbf{J}_{\mathrm{cc}}]_{i,j} - \delta_{i\,j}$ and $[\mathbf{V}_{\mathbf{eff}}]_{i,j} = \delta_{i\,j} - [\mathbf{J}_{\mathrm{cc}}]_{i,j}$
- for $i \leq N$ and $j > N$, $[\mathbf{M}_{\mathbf{eff}}]_{i,j} = [\mathbf{J}_{\mathrm{ct}}^{\mathrm{act}}]_{i,\,(j-N)}$ and $[\mathbf{V}_{\mathbf{eff}}]_{i,j} = 0$
- for $i > N$ and $j \leq N$, $[\mathbf{M}_{\mathbf{eff}}]_{i,j} = \frac{1}{\tau_t}\, [\mathbf{J}_{\mathrm{tc}}^{\mathrm{act}}]_{(i-N),\,j}$ and $[\mathbf{V}_{\mathbf{eff}}]_{i,j} = 0$
- for $i > N$ and $j > N$, $[\mathbf{M}_{\mathbf{eff}}]_{i,j} = -\frac{\delta_{i\,j}}{\tau_t}$ and $[\mathbf{V}_{\mathbf{eff}}]_{i,j} = \frac{\delta_{i\,j}}{\tau_t}$.

Note that, for conciseness, we do not explicitly add a preparatory or motif index to $\mathbf{M}_{\mathbf{eff}}$, but we warn the reader that in the following sections the matrix we refer to as $\mathbf{M}_{\mathbf{eff}}$ will be different when it is used in the context of the production of different motifs or during motif preparation.

Finally, we get:

$$\dot{\mathbf{z}} = \mathbf{M}_{\mathbf{eff}}\, \mathbf{z} \tag{S.18}$$

where $\mathbf{z} = \mathbf{z}_{\mathbf{abs}} + {\mathbf{M}_{\mathbf{eff}}}^{-1}\, \mathbf{V}_{\mathbf{eff}}\, \bar{\mathbf{z}}$ and $\left({\mathbf{M}_{\mathbf{eff}}}^{-1}\, \mathbf{V}_{\mathbf{eff}}\, \bar{\mathbf{z}}\right)$ is a bias that is fixed during a particular motif or during the preparatory period. We will refer to $\mathbf{z}$ as the 'centered rates' (representing deviations of the rates above or below some average value). We will now examine how to derive appropriate thalamocortical weight vectors and constant biases, first tackling the question of the motif dynamics and then addressing the case of the preparatory period.

**Motif dynamics** We will start by defining, for a given motif, a full vector of ideal initial conditions for the centered rates $\mathbf{z}_\mu$ of the network as a natural extension of the centered rates derived in the instantaneous thalamus framework: for $i \leq N$, $[\mathbf{z}_\mu]_i = [\mathbf{c}_\mu]_i$, and $[\mathbf{z}_\mu]_{N+1} = \mathbf{v}^\intercal\,\mathbf{c}_\mu$. Finally, we can express the dynamics of $\mathbf{z}$ as a function of the right and left eigenvectors, and the eigenvalues, of $\mathbf{M_{eff}}$ such that $\mathbf{z}(t) = \mathbf{R_{eff}}\ \mathrm{diag}\left(e^{\boldsymbol{\lambda}_{\mathbf{eff}}\,t}\right)\ \mathbf{L_{eff}}\,\mathbf{z}_\mu$.

**Tuning u to minimize the thalamic delay effects while enforcing instantaneous-loop eigenvalue control with v** Now that we expressed centered $\mathbf{z}$ dynamics, we can write an equation to ensure that, during motif production, the readout is robust to both noise in the initial conditions and to the deviations in the network response relative to the idealized immediate thalamic response. More precisely, while the idealized eigenspectrum is constrained by setting the thalamocortical weight vectors $\mathbf{v}$ through equation 8, the corticothalamic weight vectors $\mathbf{u}$ is adjusted to minimize the following cost function $C_\mathrm{m}$:

$$C_\mathrm{m} = E_{\delta\mathbf{z_0}}\left[\int_{t=0}^{t=t_\mu}\left\|\left(\mathbf{w_{eff}}^\intercal\,\mathbf{R_{eff}}\ \mathrm{diag}\left(e^{\boldsymbol{\lambda}_{\mathbf{eff}}\,t}\right)\ \mathbf{L_{eff}}\ \left(\mathbf{z}_\mu + \delta\mathbf{z_0}\right)\right) - \boldsymbol{\alpha}_{\boldsymbol{\mu}}^\intercal\, e^{\left(\tilde{\boldsymbol{\lambda}}-1\right)t}\right\|^2 dt\right], \tag{S.19}$$

where $t_\mu$ is the motif duration, $\boldsymbol{\alpha}_{\boldsymbol{\mu}}^\intercal\, e^{\left(\tilde{\boldsymbol{\lambda}}-1\right)t} = \hat{y}(t)$ is the weighted eigenmode sum which would form the readout in the idealized immediate thalamic network, and $\delta\mathbf{z_0}$ is a zero-centered and uncorrelated noise in the initial conditions with a standard deviation $\sigma_{\delta\mathbf{z_0}}$ scaling as 5% of the square root of the mean square activity norm during the motif (similarly to Eq. 11).

After a few algebraic developments, we find:

$$\begin{cases} C_\mathrm{m} = \dfrac{\mathbf{w_{eff}}^\intercal}{t_\mu}\mathbf{R_{eff}}\Bigg[\left(\left(\mathbf{L_{eff}}\left(\mathrm{diag}\left(\sigma^2_{\delta\mathbf{z_0}}\,\mathbb{1}\right) + \mathbf{z}_\mu\,\mathbf{z}_\mu^\intercal\right)\mathbf{L_{eff}}^\intercal\right)\odot\boldsymbol{\Lambda}_\mathrm{eff}\right)\mathbf{R_{eff}}^\intercal \\ \qquad - 2\left(\left(\mathbf{L_{eff}}\,\mathbf{z}_\mu\ \left(\mathbf{c}^\mathrm{E}_\mu\right)^\intercal\left(\tilde{\mathbf{L}}^\mathrm{E}\right)^\intercal\right)\odot\mathbf{B}\right)\left(\tilde{\mathbf{R}}^\mathrm{E}\right)^\intercal\Bigg]\mathbf{w_{eff}} + \dfrac{\mathbf{w}^\intercal}{t_\mu}\tilde{\mathbf{R}}\left[\left(\tilde{\mathbf{L}}\,\mathbf{c}_\mu\,\mathbf{c}_\mu^\intercal\,\tilde{\mathbf{L}}^\intercal\right)\odot\mathbf{Q}\right]\tilde{\mathbf{R}}^\intercal\mathbf{w} \\ \sigma^2_{\delta\mathbf{z_0}} = 0.05^2\ \dfrac{1}{t_\mu\,N_z}\mathbf{z}_\mu^\intercal\,\mathbf{L_{eff}}^\intercal\left[\left(\mathbf{R_{eff}}^\intercal\,\mathbf{R_{eff}}\odot\boldsymbol{\Lambda}_\mathrm{eff}\right)\right]\mathbf{L_{eff}}\,\mathbf{z}_\mu \end{cases}$$

where the twiddled letters relate to the eigendecomposition of $(\mathbf{J}_\mathrm{cc} + \mathbf{u}\mathbf{v}^\intercal)$, and the upper E index indicates an extension of arrays with a last additional zero line (initial rates vector) or last additional zero line and column (left and right eigenvector matrix). Also, we introduced $[\boldsymbol{\Lambda}_\mathrm{eff}]_{i,j} = \frac{e^{\left([\boldsymbol{\lambda}_\mathrm{eff}]_i + [\boldsymbol{\lambda}_\mathrm{eff}]_j\right)t_\mu} - 1}{[\boldsymbol{\lambda}_\mathrm{eff}]_i + [\boldsymbol{\lambda}_\mathrm{eff}]_j}$; $[\mathbf{Q}]_{i,j} = \frac{e^{\left([\tilde{\boldsymbol{\lambda}}]_i + [\tilde{\boldsymbol{\lambda}}]_j - 2\right)t_\mu} - 1}{[\tilde{\boldsymbol{\lambda}}]_i + [\tilde{\boldsymbol{\lambda}}]_j - 2}$ where $\tilde{\boldsymbol{\lambda}}$ is the vector containing the eigenvalues of $(\mathbf{J}_\mathrm{cc} + \mathbf{u}\mathbf{v}^\intercal)$; and $[\mathbf{B}]_{i,j} = \frac{e^{\left([\boldsymbol{\lambda}_\mathrm{eff}]_i + [\tilde{\boldsymbol{\lambda}}^\mathrm{E}]_j\right)t_\mu} - 1}{[\boldsymbol{\lambda}_\mathrm{eff}]_i + [\tilde{\boldsymbol{\lambda}}^\mathrm{E}]_j}$ where for $j \leq N$, $\left[\tilde{\boldsymbol{\lambda}}^\mathrm{E}\right]_j = \left(\left[\tilde{\boldsymbol{\lambda}}\right]_j - 1\right)$ and $\left[\tilde{\boldsymbol{\lambda}}^\mathrm{E}\right]_{N+1} = 0$. Finally, we also defined $N_z$ as the number of elements in $\mathbf{z}$, i.e. here $(N+1)$.

Hence, by optimizing the vector $\mathbf{u}$ to minimize the cost $C_\mathrm{m}$, while fixing the vectors $\mathbf{v}$ and $\mathbf{c}_\mu$ according to equations 8 and 5, we ensure that the effective network governing the motif dynamics is designed to ensure eigenvalue control in the limit of an infinitely fast thalamus while correcting for the deviations due to the finite thalamic timescale.

**Designing an input to the rate equations to impose positive rates during motif production** Now that we have found thalamocortical weight vectors $\mathbf{u}$ and $\mathbf{v}^\intercal$ which still perform a similar effective cortical eigenvalue control as in the main text while accounting for the thalamus' internal dynamics, we can consider the question of the biases in the rate dynamics that impose positive rates in the circuit. We will consider the noise-free non-centered rate dynamics $\mathbf{z_{abs}} = \mathbf{z} - \mathbf{M_{eff}}^{-1}\,\mathbf{V_{eff}}\,\bar{\mathbf{z}}$. Then we can write

$$\forall t, \mathbf{z_{abs}}(t) > \mathbf{0} \Leftrightarrow -\mathbf{M_{eff}}^{-1}\,\mathbf{V_{eff}}\,\bar{\mathbf{z}} > \max_t\left(-\mathbf{R_{eff}}\ \mathrm{diag}\left(e^{\boldsymbol{\lambda}_{\mathbf{eff}}\,t}\right)\ \mathbf{L_{eff}}\,\mathbf{z_0}\right) = \mathbf{b}_\mu\,.$$

We therefore set

$$\begin{aligned} -\mathbf{M_{eff}}^{-1}\,\mathbf{V_{eff}}\,\bar{\mathbf{z}} &= (\mathbf{b}_\mu + \epsilon) \Leftrightarrow \\ \bar{\mathbf{z}} &= -\,\mathbf{V_{eff}}^{-1}\,\mathbf{M_{eff}}\,(\mathbf{b}_\mu + \epsilon)\,, \end{aligned} \tag{S.20}$$

where we chose $\epsilon = 0.1$.

This will ensure that the rate variables $\mathbf{z_{abs}}$ stay positive while undergoing the dynamics in Eq. S.17 if, at the beginning of motif $\mu$, these rates are exactly set to $\mathbf{z_{abs}}_\mu = \mathbf{z}_\mu - \mathbf{M_{eff}}^{-1}\,\mathbf{V_{eff}}\,\bar{\mathbf{z}}$. However, a deviation from the right initial conditions can cause the linear rate trajectories to go beneath zero. Having a good preparatory period will therefore prove critical to ensuring that the linear dynamics solution during a motif would be very close to the solution of a rectified linear dynamical system. Fortunately, as we will now show, we can design an efficient thalamic preparatory network obeying the constraints of positive rates and non-instantaneous thalamus.

**Preparatory period dynamics** We again proceed in two steps. First, we consider the centered rates $\mathbf{z}$ with non-instantaneous thalamic dynamics (Eq. S.18). We optimize the loop weights $\mathbf{J}_{\text{tc}}^{\text{act}}$ and $\mathbf{J}_{\text{ct}}^{\text{act}}$ – where we remind the reader that the index 'act' indicates that these matrices are restricted to the subset of thalamic neurons engaged in motor preparation – to minimize a cost function $C_{\text{p}}$. Similarly to the case of instantaneous dynamics presented in the main text, $C_{\text{p}}$ integrates the distance to the steady-state centered rates $\mathbf{z_{ss}}$:

$$\begin{aligned} C_{\text{p}} &= E_{\mathbf{z_0}}\left[\int_0^\infty \|\mathbf{z}(\mathbf{t}) - \mathbf{z_{ss}}\|^2\,dt\right] \\ &= \sigma_{\mathbf{z_0}}\ \mathrm{Tr}\left(\mathbf{R_{eff}}\left((\mathbf{L_{eff}}\,\mathbf{L_{eff}}^{\mathsf{T}}) \odot \mathbf{A}\right)\mathbf{R_{eff}}^{\mathsf{T}}\right) \end{aligned}$$

where $[\mathbf{A}]_{i,j} = \frac{-1}{[\boldsymbol{\lambda}_{\text{eff}}]_i + [\boldsymbol{\lambda}_{\text{eff}}]_j}$ and $\sigma_{\mathbf{z_0}}$ is the standard deviation of the random initial conditions $\mathbf{z_0}$ which are assumed to be centered and iid.

We then consider which biases $\bar{\mathbf{z}}_{\mathbf{prep}}$ to add to the preparatory dynamics (Supplementary Eq. S.17) to get positive rates $\mathbf{z_{abs}} = \mathbf{z} - \mathbf{M_{eff}}^{-1}\,\mathbf{V_{eff}}\,\bar{\mathbf{z}}_{\mathbf{prep}}$. We remind the reader that $\mathbf{M_{eff}}$ is now a preparatory-period specific effective connectivity, which is different from the motif-specific connectivities considered in the previous paragraph though the notation is the same for conciseness.

The difficulty is to find biases that are sufficient to create positive rates in the thalamocortical preparatory network despite the variability of trajectories among different transitions. On the one hand, we can use the same method as in the main text to impose that cortical rates converge towards the positive activity pattern which is appropriate to start the next motif (main text section 4.4), hence ensuring rate positivity at the end of the preparatory period. On the other hand, this method does not give guidelines for the inputs needed in thalamus, and cannot by itself impose positive rates at the beginning of the preparatory period.

Therefore, next, we first focus on estimating average biases that can ensure positive rates within the preparatory network regardless of the variability of initial rates at the beginning of the motif transition period (which reflects the rate variability when producing the preceding motifs). After this, we will use these average biases to craft upcoming-motif-specific biases which will both ensure the positivity of the rates and the convergence of the cortical rates towards the appropriate pattern of activity at the end of the preparatory period.

**Average biases able to enforce rate positivity during the preparation period despite initial rates variability** Here, we treat the absolute rates at the beginning of the preparatory period $\mathbf{z}_{\mathbf{abs\,0}}^{\mathbf{prep}}$ as an i.i.d. random vector with mean $\boldsymbol{\mu}_{\mathbf{z_{abs\,0}}}^{\mathbf{prep}}$ and standard deviation $\boldsymbol{\sigma}_{\mathbf{z_{abs\,0}}}^{\mathbf{prep}}$. For the cortical units, the pattern of activity at the beginning of the preparatory period corresponds to the pattern of activity at the end of motifs. Hence, we simply estimate $\{[\boldsymbol{\mu}_{\mathbf{z_{abs\,0}}}^{\mathbf{prep}}]_i\,,\ [\boldsymbol{\sigma}_{\mathbf{z_{abs\,0}}}^{\mathbf{prep}}]_i\}_{i\leq N}$ by computing the mean and standard deviation of $\mathbf{z}$ over all units and all motifs at the end of six example motifs (the sinc function as in Fig. 2 and slow oscillating motifs as in Fig. 4). In addition, for the statistics of the rates in the preparatory network units: $\{[\boldsymbol{\mu}_{\mathbf{z_{abs\,0}}}^{\mathbf{prep}}]_i\,,\ [\boldsymbol{\sigma}_{\mathbf{z_{abs\,0}}}^{\mathbf{prep}}]_i\}_{i>N}$, we use a very small positive value for both the mean and the standard deviation to account for the fast rate decay of these unconnected neurons in between two preparatory periods when they are unconnected from cortex.

We then design a cost function $C_{\mathrm{p}}^{\mathrm{abs}}$ to find bias terms $\mathbf{b_{eff}^{prep}} = -\mathbf{M_{eff}}^{-1}\,\mathbf{V_{eff}}\,\bar{\mathbf{z}}^{\mathbf{prep}}$ such that the random variable:

$$\mathbf{z_{abs}^{prep}}(t) = \mathbf{R_{eff}}\ \mathrm{diag}\left(e^{\boldsymbol{\lambda}_{\mathrm{eff}}\,t}\right)\ \mathbf{L_{eff}}\ \mathbf{z_0^{prep}} + \mathbf{b_{eff}^{prep}} \tag{S.21}$$

is likely to be positive at all times during the preparatory period. The stochasticity of $\mathbf{z_{abs}^{prep}}(t)$ emerges because $\mathbf{z_0^{prep}} = \mathbf{z_{abs\,0}^{prep}} - \mathbf{b_{eff}^{prep}}$ is a random vector with mean $\boldsymbol{\mu}_{\mathbf{z_0}}^{\mathbf{prep}} = \boldsymbol{\mu}_{\mathbf{z_{abs\,0}}}^{\mathbf{prep}} - \mathbf{b_{eff}^{prep}}$ and standard deviation $\boldsymbol{\sigma}_{\mathbf{z_0}}^{\mathbf{prep}} = \boldsymbol{\sigma}_{\mathbf{z_{abs\,0}}}^{\mathbf{prep}}$. As a step to define the cost function $C_{\mathrm{p}}^{\mathrm{abs}}$, we can express the statistics of $\mathbf{z_{abs}^{prep}}(t)$ at all times to find the vector of trajectories $\mathbf{h_{low}}(\mathbf{t})$ which we define as the value taken at three standard deviations below the mean for each unit:

$$\begin{cases} E_{\mathbf{z_0^{prep}}}\left[\mathbf{z^{prep}}(t)\right] = \mathbf{R_{eff}}\ \mathrm{diag}\left(e^{\boldsymbol{\lambda}_{\mathrm{eff}}\,t}\right)\ \mathbf{L_{eff}}\ \left(\boldsymbol{\mu}_{\mathbf{z_{abs\,0}}}^{\mathbf{prep}} - \mathbf{b_{eff}^{prep}}\right) = \mathbf{f}(\mathbf{t}) \\ E_{\mathbf{z_0^{prep}}}\left[\mathbf{z^{prep}}(t) - E_{\mathbf{z_0^{prep}}}\left[\mathbf{z^{prep}}(t)\right]\right]^2 = \\ \quad \mathbf{R_{eff}}\ \mathrm{diag}\left(e^{\boldsymbol{\lambda}_{\mathrm{eff}}\,t}\right)\ \mathbf{L_{eff}}\ \mathrm{diag}\left(\left(\boldsymbol{\sigma}_{\mathbf{z_{abs\,0}}}^{\mathbf{prep}}\right)^2\right)\ \mathbf{L_{eff}}^{\intercal}\ \mathrm{diag}\left(e^{\boldsymbol{\lambda}_{\mathrm{eff}}\,t}\right)\ \mathbf{R_{eff}}^{\intercal} = \mathbf{g}(\mathbf{t}) \\ \mathbf{h_{low}}(\mathbf{t}) = E_{\mathbf{z_{abs\,0}^{prep}}}\left[\mathbf{z_{abs}^{prep}}(t)\right] - 3 * \sqrt{E_{\mathbf{z_{abs\,0}^{prep}}}\left[\mathbf{z_{abs}^{prep}}(t) - E_{\mathbf{z_{abs\,0}^{prep}}}\left[\mathbf{z_{abs}^{prep}}(t)\right]\right]^2} = \mathbf{b_{eff}^{prep}} + \mathbf{f}(\mathbf{t}) - 3 * \sqrt{\mathbf{g}(\mathbf{t})} \end{cases}$$

To ensure that, under linear dynamics during the preparatory period, the rates stay as positive as possible, we therefore optimize $\mathbf{b_{eff}^{prep}}$ to minimize the sum of the square of the negative values of $\min_t\left(\mathbf{h_{low}}(\mathbf{t})\right) = \mathbf{h_{low}^{min}}$:

$$C_{\mathrm{p}}^{\mathrm{abs}} = \sum_{i\ s.t.\ \mathbf{h_{low}^{min}}(i)<0} \left(\mathbf{h_{low}^{min}}(i)\right)^2$$

where $\mathbf{h_{low}}(\mathbf{t})$ was minimized over a preparatory period duration of five cortical timescales.

Finally, we remark that in this scenario where we managed to force the rates $\mathbf{z_{abs}^{prep}}$ to be mostly positive at all times, the rates converge towards $\mathbf{b_{eff}^{prep}}$, which means that some more work is needed to ensure that during the preparatory period the rates not only stay positive but also converge the appropriate cortical activity pattern to start the next motif – which is the topic of the next section.

**Biases ensuring rate positivity during the preparation towards a specific upcoming motif** The biases $\mathbf{b_{eff}^{prep}} = -\mathbf{M_{eff}}^{-1}\,\mathbf{V_{eff}}\,\bar{\mathbf{z}}_{\mathbf{prep}}$ are designed such that under linear dynamics $\dot{\mathbf{z}}_{\mathbf{abs}}^{\mathbf{prep}} = \mathbf{M_{eff}}\,\mathbf{z_{abs}^{prep}} + \mathbf{V_{eff}}\,\bar{\mathbf{z}}^{\mathbf{prep}}$, the rates are likely to be positive within the preparatory network, despite the variability of the initial rates among different preceding motifs.

In addition, we need to ensure that at the end of the preparatory period, the positive cortical rates converge towards the desired positive pattern to start the next motif $\mu$: $\mathbf{z}_{\mathbf{abs}\,\mu} = \mathbf{z}_\mu + (\mathbf{b}_\mu + \epsilon)$ (as defined in Eq. S.20). Similar to the methods used in the main text, it is easy to see that this relies on designing an additive input $\bar{\mathbf{z}}_\mu^{\mathbf{prep}}$ specific to the dynamics leading to a particular motif:

$$\dot{\mathbf{z}}_{\mathbf{abs}}^{\mathbf{prep}} = \mathbf{M_{eff}}\ \mathbf{z_{abs}^{prep}} + \mathbf{V_{eff}}\ \bar{\mathbf{z}}_\mu^{\mathbf{prep}} \Leftrightarrow \lim_{t\to\infty} \mathbf{z_{abs}^{prep}}(t) = -\mathbf{M_{eff}}^{-1}\ \mathbf{V_{eff}}\ \bar{\mathbf{z}}_\mu^{\mathbf{prep}} = \mathbf{z}_{\mathbf{abs}\,\mu} \tag{S.22}$$

The cortical steady-state rates are constrained by the ideal pattern to start the upcoming motif. We however still have some freedom for choosing the steady-state thalamic preparatory rate. This is where we use the average positive dynamics $(\mathbf{z_{prep}} + \mathbf{b_{eff}^{prep}})$ (defined in Eq. S.21), which converge towards $\mathbf{b_{eff}^{prep}}$ and maintain rate positivity despite the initial variability in the rates among different transitions, to determine the thalamic steady-state rates which can ensure positive dynamics within the thalamocortical circuit even in the presence of the relevant rate variability. Hence, we set, for $i \leq N$, $\left[\mathbf{z}_{\mathbf{abs}\,\mu}\right]_i = \left[\mathbf{c}_\mu\right]_i + \left[\mathbf{b}_\mu + \epsilon\right]_i$; else, for $i > N$, $\left[\mathbf{z}_{\mathbf{abs}\,\mu}\right]_i = \frac{\sum_{j=N+1}^{N\,+\,N_{\text{prep. thal.}}}\left[\mathbf{b_{eff}^{prep}}\right]_j}{N_{\text{prep. thal.}}}$, where $N_{\text{prep. thal.}}$ is the number of units in the preparatory network.

Hence, for each transition towards a particular motif, we determined a vector $\bar{\mathbf{z}}_{\mu}^{\mathbf{prep}} = -\mathbf{V_{eff}}^{-1}\,\mathbf{M_{eff}}\,\mathbf{z_{abs}}_{\mu}$ such that we can write an efficient and mostly positive preparatory dynamics according to equation S.22. In Supplementary Fig. S.6b, we show the success of this approach on thirty-six example transitions (all possible transitions among the sinc function as in Fig. 2 and five slow oscillating motifs as in Fig. 4).

**Producing and reading out a motif sequence in network with positive rates and realistic thalamic time constant** To produce sequential dynamics, we can string together the motif-specific and preparatory dynamics described above, obeying Supplementary Eq. S.16.

We emphasize that, in Supplementary Eq. S.16, $\bar{\mathbf{r}}$ and $\overline{\mathbf{t_{act}}}$ are biases that are specific to the stage of the sequence that the network is producing: either the motif currently produced or the upcoming motif during the preparatory period; in addition, the index 'act' indicates the parts of the thalamocortical circuit that are specific either to the motif being currently produced or to the preparatory period.

To assess the effectiveness of the biases in keeping the rates positive, we compare the linear dynamics described in Supplementary Eq. S.16 to rectified non-linear dynamics:

$$\begin{cases} \dot{\mathbf{c_{abs}}} = -\left(\mathbf{c_{abs}} - \bar{\mathbf{c}}\right) + \mathbf{J}_{\text{cc}}\left(\left[\mathbf{c_{abs}}\right]^{+} - \bar{\mathbf{c}}\right) + \mathbf{J}_{\text{ct}}^{\text{act}}\ \left[\mathbf{t_{act}}\right]^{+} \\ \dot{\mathbf{t_{act}}} = \dfrac{1}{\tau_t}\left(-\left(\mathbf{t_{act}} - \overline{\mathbf{t_{act}}}\right) + \mathbf{J}_{\text{tc}}^{\text{act}}\ \left[\mathbf{c_{abs}}\right]^{+}\right) \end{cases} \tag{S.23}$$

Finally, for both the linear and rectified dynamics, we also have to think about the effect of the biases on the readout. For any motif, adding the biases $\bar{\mathbf{c}}$ and $\overline{\mathbf{t_{act}}}$ –regrouped into the vector $\bar{\mathbf{z}}$– to the differential equations transforms the centered dynamics $\mathbf{z}$, which is crafted to match the desired output, into the offset dynamics $\mathbf{z_{abs}} = \mathbf{z} - \mathbf{M_{eff}}^{-1}\,\mathbf{V_{eff}}\,\bar{\mathbf{z}}$.

Hence, to recover the desired readout during a particular motif, we need to remove the motif-dependent constant $\mathbf{w_{eff}}\ \mathbf{M_{eff}}^{-1}\ \mathbf{V_{eff}}\ \bar{\mathbf{z}}$ from the readout, where the effective readout weights $\mathbf{w_{eff}}$ are defined such that for $i \leq N$ $[\mathbf{w_{eff}}]_i = [\mathbf{w}]_i$, else $[\mathbf{w_{eff}}]_i = 0$. In addition, during the transitions between motifs, we choose to favor the continuity of the readout with the upcoming motif. To this aim, we feed the readout the with same constant as for the upcoming motif during the preparatory period.

Note that this added input to the readout could potentially be avoided if the eigenvalue control would include, in addition to eigenmodes for fitting the desired motifs, an additional eigenvalue close to zero real and imaginary part, which would allow to have the additional freedom of producing a constant value during the readout.

#### 5.2 Supplementary figures and table

##### 5.2.1 Table summarizing the assumptions of the model and their justification

In view of available evidence, we make series of assumptions to constrain and simplify our corticothalamic model and summarize those assumptions in the following table. Lines 1–2 concern the modeling of cortical population rate dynamics as a dynamical system with quasi-linear motif-specific dynamics with a quasi-linear mapping from cortex to muscle activation. Lines 3–4 concern the anatomy of the cortex–thalamus complex and its dynamical consequences. Lines 5–7 concern several assumptions for the thalamic dynamics and the role of basal ganglia input to thalamus.

| | Assumption | Source of evidence | References |
|---|---|---|---|
| 1 | Linear readout from recurrent motor cortical population dynamics as an approximate motor output | Anatomy / Electrophysiology | [48, 26, 4, 6, 80] |
| 2 | Quasi-linear input–output relation in balanced neural populations; fit of motor cortical population activity with linear dynamics during individual behaviors | Theory / Electrophysiology | [89, 90, 79] [6] |
| 3 | Non-recurrent thalamus bidirectionally connected to cortex | Anatomy | [54, 49] |
| 4 | Slower cortical dynamics (relative to thalamus) because of cortical excitatory recurrence (absent in the thalamus) | Electrophysiology / Theory | [83, 72, 48, 6] |
| 5 | Quasi-rectified linear tonic firing in thalamus | Electrophysiology | [85, 61] but see [27] |
| 6 | Strong inhibitory connections from basal ganglia to thalamus | Electrophysiology / Optogenetics | [9, 10, 27], but see [47] |
| 7 | Sequence-element related activity in basal ganglia (sustained during motifs and phasic at switch times) | Neurophysiology | [24, 25] |

Table 1: Summary of model assumptions and their justifications

##### 5.2.2 Limits of eigenvalue control through a single thalamocortical loop

Here we tackle in more detail the question of the existence and numerical stability of solutions for the control of eigenvalues using equation 8.

In the limiting case when the number $K$ of target eigenvalues $\lambda^*$ matches the number of cortical neurons $N$, the pseudoinverse can be replaced by a simple matrix inverse, in which case the question of the existence of a solution to the eigenvalue control problem boils down to the invertibility of the matrix $\mathbf{P}$. If the initial matrix $\mathbf{J}_{\mathrm{cc}}$ is random, full-rank and without degeneracies such that all $\lambda_i$ are distinct and random, and if the target eigenvalues $\lambda_i^*$ are also distinct from each other and from the initial eigenspectrum, then the matrix $\mathbf{P}$ with elements $P_{ij} = (\lambda_i^* - \lambda_j)^{-1}$ is also almost certainly invertible. However, $\mathbf{P}$ might be more or less well-conditioned such that the inversion is more or less numerically stable. If there are several target eigenvalues outside of the initial eigenspectrum, $\mathbf{P}$ will tend to have correlated rows and may therefore be difficult to invert. Accordingly, we found that full eigenvalue control using Eq. 8 was numerically feasible if the target eigenvalues were chosen from the same distribution as the eigenvalues of $\mathbf{J}_{\mathrm{cc}}$ (Supplementary Fig. S.1b), while it failed when the target eigenvalues were chosen from a different partially-overlapping distribution (Supplementary Fig. S.1c). In addition, in the latter case, the (numerically inaccurate) solution involved very large weights [88] (Supplementary Fig. S.1g, compare to panels e and f). These issues can be mitigated if only choosing a few target eigenvalues $\lambda_i^*$ such that eigenvalue control only requires solving an underdetermined linear system using the pseudoinverse of $\mathbf{P}$ (Supplementary Fig. S.1d and h).

##### 5.2.3 Eigenmodes as an efficient set of basis functions to fit transients

For each motif, our goal is to match the output of the network $\hat{y} = \mathbf{w}^\intercal \mathbf{c}$ with a desired weighted sum of a few complex conjugate exponentials $y^* = \sum_{i=1}^{K} \alpha_i e^{(\lambda_i^*-1)t}$ (see the dashed lines in Fig. 1e). Here we illustrate that complex conjugate exponentials – where each pair sums to a periodic function that is exponentially modulated and real – constitute a useful set of basis functions which can be efficiently linearly combined to approximate many target motifs $y$ (as was argued based on theoretical arguments in [55]). More specifically,

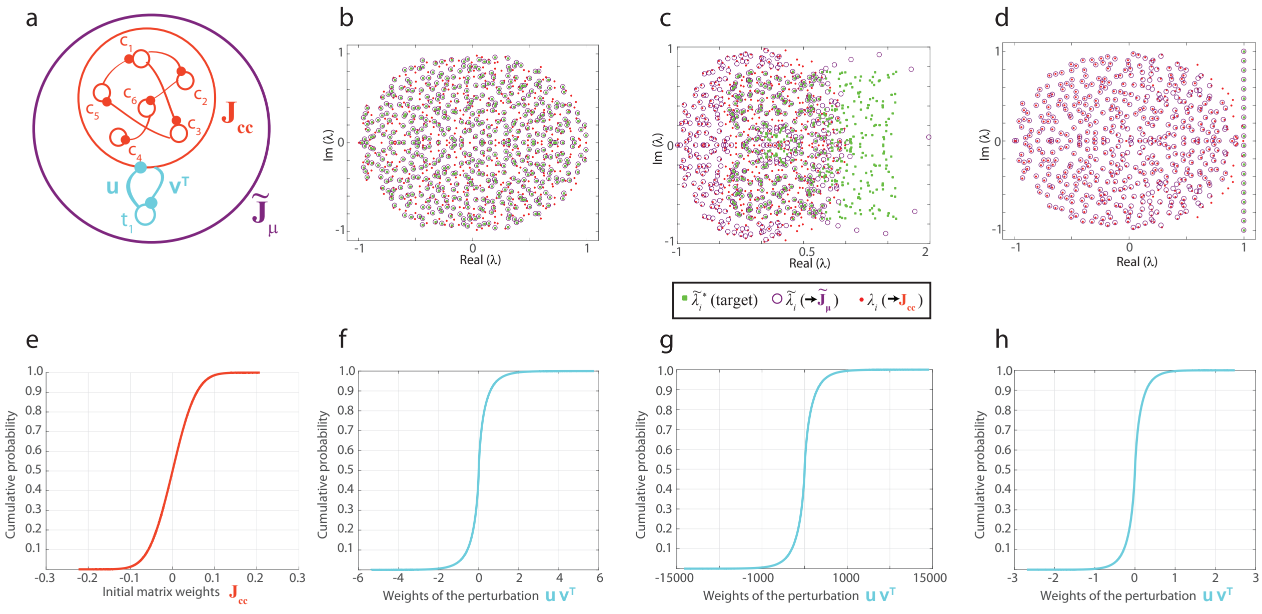


Figure S.1: **a)** The cortex (with recurrent weights $\mathbf{J}_{cc}$ and eigenvalues $\lambda_\mu$) interacts with one thalamocortical loop $\mathbf{u}\mathbf{v}^\intercal$, creating an effective recurrent network with connectivity $\tilde{\mathbf{J}}_\mu = \mathbf{J}_{cc} + \mathbf{u}\mathbf{v}^\intercal$ and eigenvalues $\tilde{\lambda}_\mu$. **b–d)** Eigenspectra of the effective connectivity matrices reached by computing the vector **v** according to Eq. 8 ($\tilde{\lambda}_i$, purple circles) compared with the desired eigenvalues $\lambda_i^*$ (green dots). The eigenspectrum of the isolated cortical connectivity matrix $\mathbf{J}_{cc}$ is given for comparision ($\lambda_i$, red dots). **b)** The $\lambda_i^*$ are the eigenvalues of a different random matrix from $\mathbf{J}_{cc}$ but with the same distribution of entries. The matrix **P** from Eq. 8 is easy to invert, so the solution for the vector **v** is relatively accurate, and the eigenvalues of the effective connectivity matrix match the desired eigenvalues. **c)** The desired eigenvalues $\lambda_i^*$ are very different from the cortical eigenvalues. The matrix **P** from Eq. 8 is now very hard to invert, and numerical imprecisions lead to a mismatch between $\lambda_i^*$ and $\tilde{\lambda}_i$. **d)** We specify only 20 desired $\lambda_i^*$ outside of the spectrum of the isolated matrix, and find a robust solution for **v** such that the effective matrix has eigenvalues that match the desired values. **e)** Cumulative probability distribution for the weights of the cortical matrix $\mathbf{J}_{cc}$. The weights come from a normal distribution with variance $1/N$. **f,g,h)** Cumulative probability distributions for the weights of the perturbation $\mathbf{u}\mathbf{v}^\intercal$ after finding **v** with Eq. 8. The perturbation weights are larger than the initial cortical weights, but less so when the change in eigenspectrum is not too pronounced. For all panels, the entries of the vector **u** are taken at random from a Gaussian distribution.

we can demonstrate that the approximation quickly improves when increasing $K$, the number of basis functions, for a variety of motif shapes. We can find the coefficients $\alpha_i$ and the timescales $\lambda_i^*$ either using numerical methods (Supplementary Fig. 1a-d) or analytics (Supplementary Fig. 1e). For a diversity of motifs, a few complex conjugate exponentials ($K = 4$–$20$, corresponding to 2–10 real exponentially modulated sine waves) are sufficient for very good fits (Supplementary Fig. 1a-b). This is true even for discontinuous motifs which are hard to approximate as they require both slow exponential decay timescales and fast oscillatory frequencies, corresponding to outlier eigenvalues far from the initial spectrum (see Supplementary Fig. S.1). For some analytic functions, such as the sinc function (i.e.,sinc$(t) = \sin(t)/t = \frac{1}{2}\int_{-1}^{1} \cos(\omega t)\, d\omega$), we can deduce from the equation a good discrete set of parameters for the exponentials. (In this case, a uniform tiling of $\lambda_i^*$ on the interval $[1 - 1i, 1 + 1i]$ with amplitudes $\alpha_i = 1/K$; Supplementary Fig. 1e).) Note that while the latter approach does not require any numerics and favors parameter values that maximize the similarity between the sum of complex exponentials and the full domain of definition of the target function, it can lead to a less efficient approximation compared to the numerical fit of a finite-time transient with a sum of a few modes.

We remark that for all the targets that we consider, the modes with eigenvalues close to the line $\mathrm{Re}\,\lambda = 1$ (quasi-pure sines) play an important role, as is reminiscent of Fourier series.

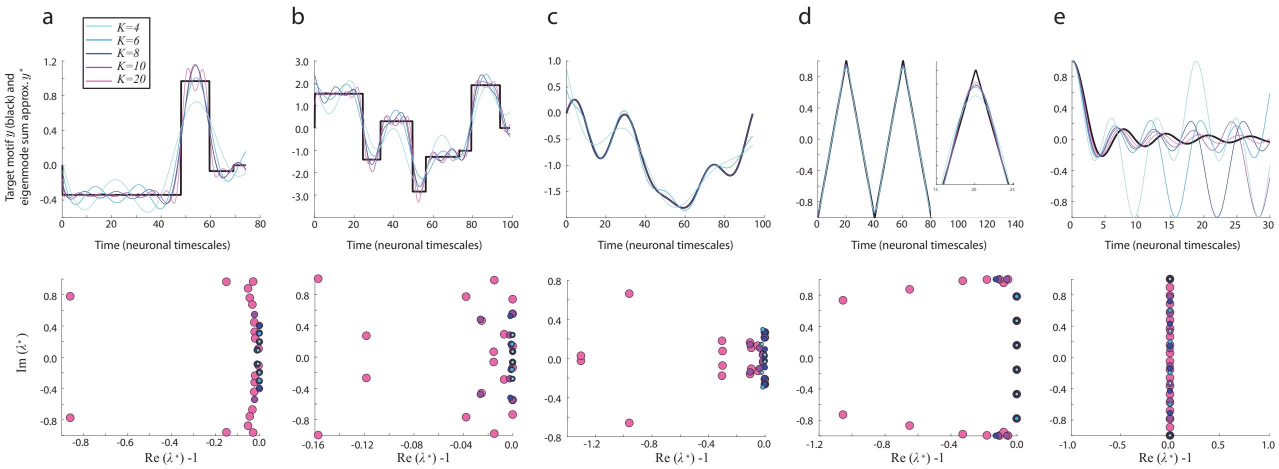


Figure S.2: **Top:** Example target functions $y$ (black) and their approximation by linear combinations of eigenmodes which are conjugate pairs of complex exponentials $y^* = \sum_{i=1}^{K} \alpha_i e^{(\lambda_i^* - 1)t}$ (from light blue to pink, $K = 4$, $K = 6$, $K = 8$, $K = 10$ and $K = 20$). **Bottom:** Real and imaginary parts of the parameters $\lambda_i^* - 1, \ldots, \lambda_N^* - 1$ used to match $y^*$ and $y$, separately for different values of $K$. Note that increasingly large symbols were used for larger values of $K$, in order to make all symbols visible in cases when the $\lambda_i^*$ overlap for different values of $K$. **a–d)** The parameters $\alpha_i$ and $\lambda_i^*$ are numerically chosen to minimize the mean square error between $y$ and $y^*$ (see Methods section 4.5 for detail). Note that in **c**, the curves for the larger values of $K$ are hidden under the $K = 8$ curve. In **d**, the inset shows a magnified part of the first upper triangle. **e)** The sinc function ($\text{sinc}(t) = \sin(t)/t = \frac{1}{2}\int_{-1}^{1} \cos(\omega t)\, d\omega$), and analytic approximations to the sinc function using discretized parameters $\lambda_i^*$ uniformly distributed between $1 - 1i$ and $1 + 1i$.

##### 5.2.4 More details on harnessing non-normal amplification in a rank-one perturbed network

Here, we show additional data concerning the emergence of eigenvector correlations and non-normal amplification in the rank-one perturbed network with effective connectivity $\tilde{\mathbf{J}}_\mu = \mathbf{J}_{cc} + \mathbf{u}\mathbf{v}^\intercal$.

**Eigenvector correlations in rank-one perturbed networks** In Fig. 2g we showed the presence of eigenvector correlations in a random recurrent network interacting with a loop designed to shape the network output into an approximation of the sinc function (Fig. 2a,d). Here, in Supplementary Fig. S.3a,b, we show the full distribution of cosine of eigenvector angles in the effective matrix $\tilde{\mathbf{J}}_\mu$, with a half-tuned or fully-tuned rank-one perturbation respectively. We defined the cosine of the angle between eigenvectors $\tilde{\mathbf{l}}_i$ and $\tilde{\mathbf{l}}_j$ as $\cos\theta_{ij} = \text{Re}\left(\tilde{\mathbf{l}}_i^\intercal \overline{\tilde{\mathbf{l}}_j}\right) / \left(\|\tilde{\mathbf{l}}_i^\intercal\| \, \|\overline{\tilde{\mathbf{l}}_j}\|\right)$. For this figure, more specifically, we computed the cosine angle after normalizing eigenvectors such that the norm of their real and imaginary parts are both 0.5 and sum to a total norm of 1. A large cosine of angle indicates a correlation (in other words, a quasi-parallelism) between eigenvectors.

Our analytic expressions for eigenvectors of $\tilde{\mathbf{J}}_\mu$ (Supplementary Sec. 5.1.2) suggests that there will be strong correlations between eigenvectors associated with eigenvalues that are relatively more similar with one another compared to the usual similarity between eigenvalues of the initial spectrum of $\mathbf{J}_{cc}$. Here, as the rank-one perturbation is used to impose outlier eigenvalues at the periphery of the initial spectrum to create a desired output (Supplementary Fig. S.3e), we expect large correlations between eigenvectors corresponding to controlled eigenvalues. To investigate whether our simulations are consistent with our theoretical predictions, we ranked the eigenvectors such that the eigenvector corresponding to the controlled eigenvalues, that are situated beyond the rightmost border of the bulk of the eigenspectrum, correspond to the upper left corner of the correlation matrix (outlined by a pink asterix and arrow), while the remaining vectors are ranked by the norm of their associated egienvalues (i.e., the vectors just after the pink arrow correspond to the eigenvalues with norm close to 0 that are at the center of the bulk of spectrum of $\tilde{\mathbf{J}}_\mu$). In agreement with our theoretical predictions, we observe large magnitudes within the top left corner of the correlation matrices in Supplementary Fig. S.3a–b – corresponding to controlled eigenvalues – but small magnitudes (cyan color) in

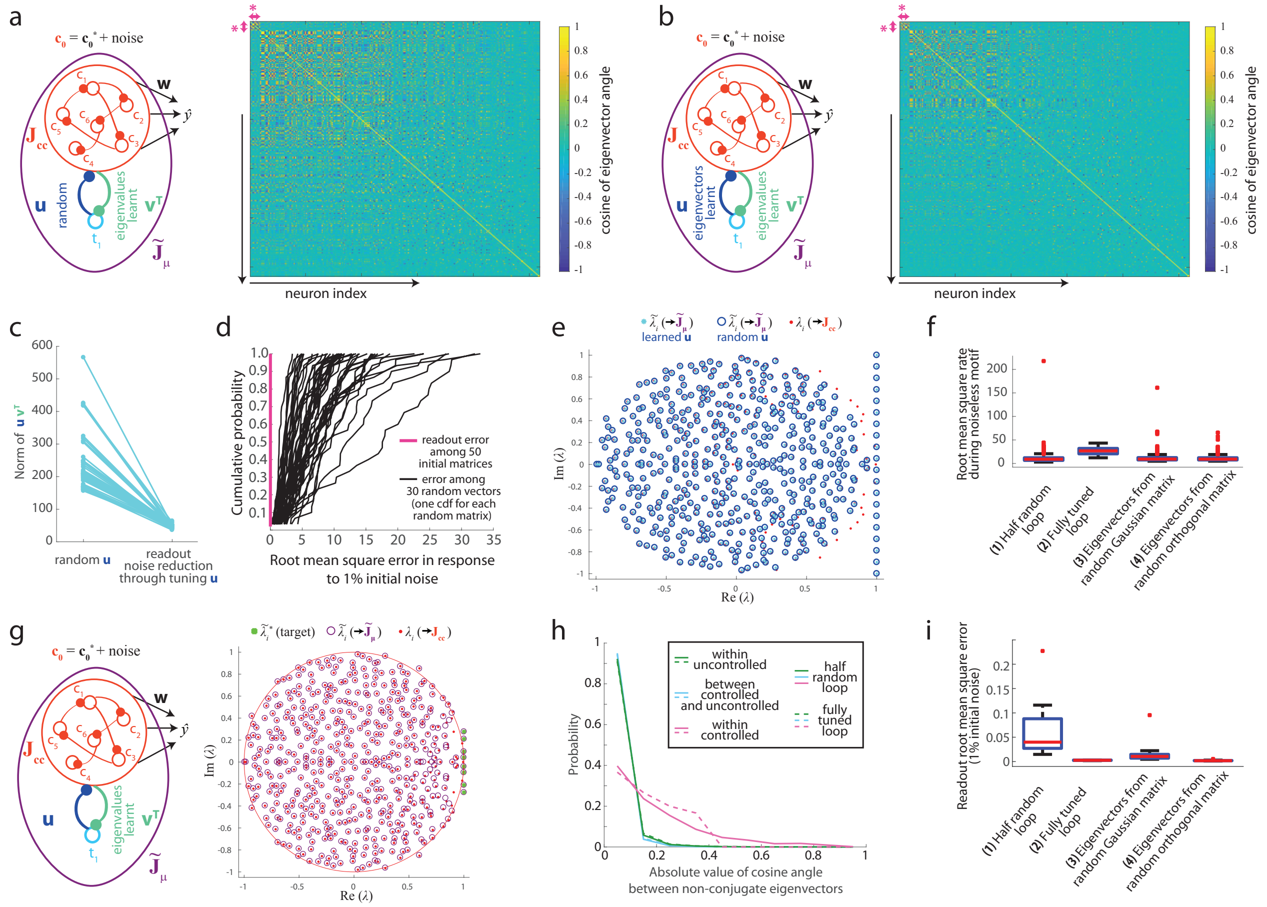


Figure S.3: **a-f)** Networks tuned for the production of the sinc function (main text Figs. 1d and 2). **a)** For a half-random thalamocortical perturbation (**left**), cosine of angle between pairs of non-complex conjugate eigenvector (**right**). Bright yellow and deep blue indicate larger correlations. Eigenvectors are ordered such that the first ten, as outlined by the pink stars, are corresponding to the controlled eigenvalues, while subsequent indices correspond to non-controlled eigenvalues ordered from smaller to larger eigenvalue norm. **b)** Same as **a** but in the case of a fully learned loop. **e)** Decrease of the Frobenius norm of $\mathbf{u}\mathbf{v}^\mathsf{T}$ between half-random and fully-tuned loops (50 different $\mathbf{J}_{cc}$ and $\mathbf{w}$). **d)** Cumulative distributions of mean squared deviation due to initial noise in networks with fully tuned loops, separately along the readout direction (pink) or along thirty different random directions (black, each line is the distribution over these random directions for a given $J_{cc}$ and $\mathbf{w}$). **e)** Eigenspectrum of $J_{cc}$ (red), and of the effective $\tilde{\mathbf{J}}_\mu$ in case of a half random perturbation(blue circles) or a fully tuned loop (cyan filled dots). **f)** Root mean square rate in networks with a half-random loop (1), after full optimization of the loop (2), or in matrices with the same eigenspectrum as $\tilde{\mathbf{J}}_\mu$ but with either eigenvectors from random matrices (3) or orthogonal eigenvectors (4). **g)** For a thalamocortical loop tuned to produce the motif 'C' from Fig. 4, eigenspectrum of the effective matrix $\tilde{\mathbf{J}}_\mu$ (purple circles) and target desired eigenvalues (green filled dots). The eigenvalues of the isolated cortical matrix $\mathbf{J}_{cc}$ are also shown (red dots) as well as the limits of the eigenspectrum that $J_{cc}$ would have if its size was infinite (red circle). **h,i)** Properties of networks optimized for five oscillatory motifs (including the three motifs of Fig. 4). **h)** Distributions of the absolute values of the cosines between eigenvectors, separately if *(i)* both eigenvectors correspond to controlled eigenvalues (magenta), *(ii)* only one of the two eigenvectors of the pair correspond to a controlled eigenvalue (cyan), *(iii)* both eigenvectors correspond to uncontrolled eigenvalues (green). We further differentiate between a half-random loop (only optimized for eigenvalue control) and a fully trained loop (also optimized for robustness). **i)** Root mean square error of the output in the presence of 1% noise. Groups defined as in **f**.

the remaining entries of the first $K$ rows and columns – indicating relatively lower correlations between the eigenvectors of controlled eigenvalues and the eigenvectors associated with the $N-K$ remaining eigenvalues. In addition, we see relatively larger correlation values in between eigenvectors with small-norm eigenvalues (upper left diagonal after the pink-arrow highlighted area), which is also as expected from our analytics as those eigenvalues are relatively clustered together compared to the rest of the eigenspectrum.

Also, in Supplementary Fig. S.3h we confirm our results from Fig. 2g showing large correlations between eigenvectors of controlled eigenvalues. For this panel, we combine the results from five motifs created from the low-pass filtering of the natural oscillations of a random chaotic network, three of which are shown in Fig. 4 and another is shown in Supplementary Fig. S.2c). These motifs can be produced with a smaller modification of the eigenspectrum of $\mathbf{J}_{cc}$ than the sinc function (compare the results of eigenvalue control in Supplementary Fig. S.3e and in Supplementary Fig. S.3g). In Supplementary Fig. S.3i, we show that even this more subtle perturbation of $\mathbf{J}_{cc}$ with a half-random perturbation leads to larger output noise than control matrices with little non-normal amplification, even though the effect is smaller than in the case of the sinc function. Finally, Supplementary Fig. S.3i also shows that a full tuning procedure of the perturbation weights similarly permits the improvement of the output's noise robustness in the case of these slow oscillatory motifs. Hence, qualitatively, the results we describe in the main text about eigenvector correlations and their functional implications in rank-one perturbed networks generalize to different types of desired output.

**Decrease of the perturbation norm between half-random and fully-tuned control procedures** From the comparison between Fig. 2c and f, we showed for one random sample of $\mathbf{J}_{cc}$ and $\mathbf{w}$ how the distribution of perturbation weights tightens in the case of a fully-tuned perturbation controlling both the eigenvalues and the eigenvectors of the circuit, compared to the case of a half-tuned perturbation controlling only eigenvalues. Here, in Supplementary Fig. S.3c, we show that this decrease of the perturbation norm following the full-tuning procedure is true across many different random samples of $\mathbf{J}_{cc}$ and $\mathbf{w}$. Note also how, after the full-tuning procedure, the variance of the perturbation norm decreases, emphasizing how this procedure takes advantage of the remaining degrees of freedom in the weight distribution.

**Anisotropy of noise amplification in fully tuned rank-one perturbed networks** From the main text Fig. 2h and i, we can indirectly conclude that in a network with a fully tuned perturbation (i.e., also optimized to minimize the effect of initial noise on the network output), the directions of noise amplification in the network are orthogonalized relative to the readout direction. Here, in Supplementary Fig. S.3c, we show this more directly by comparing the magnitude of the effect of initial deviations between *(i)* the timecourse of the activities projected on random vectors and *(ii)* the output of the network, over different randomly drawn vectors, cortical matrices $\mathbf{J}_{cc}$, and readout weights $\mathbf{w}$. The deviations are indeed significantly larger along random vectors than along the readout direction.

**Conservation of the full eigenspectrum between half-random and fully-tuned rank-one perturbations** In Fig.1e, we showed the eigenspectrum of an effective connectivity matrix including a half-random rank-one perturbation to impose a few target eigenvalues $\lambda_1^*, \ldots, \lambda_K^*$. Here, in Supplementary Fig. S.3e, we show that besides the $K$ chosen eigenvalues, the other $N-K$ eigenvalues are also conserved between any arbitrary half-random perturbation (random $\mathbf{u}$ and $\mathbf{v}$ chosen using equation 8) and a fully-learned perturbation ($\mathbf{u}$ tuned to minimize the effect of initial noise on the output with $\mathbf{v}$ chosen using equation 8). This emphasizes that the optimization of $\mathbf{u}$ to improve output noise robustness only relies on tuning the eigenvectors of the effective connectivity matrix, in agreement with our analytical results (see equation S.2).

**Norm of the neuronal activities in rank-one perturbed network** In Fig. 2e, we suggest that in the effective network with a fully-tuned rank-one perturbation, though the directions of large noise amplification are orthogonalized with respect to the readout $\mathbf{w}$, the eigenvectors associated with (the slow) controlled eigenvalues stay relatively aligned with $\mathbf{w}$. This is because if this was not the case, then the activity norm in the network would need to be very large in order to get an output with non-zero amplitude, as is clear from equation 5. Though this configuration could potentially appear noise robust if assuming a noise of fixed amplitude, which would therefore become relatively smaller as the activity norm grows, this is not a desirable outcome of noise robustness optimization. Indeed, very large activity in a network is unrealistic, and the neuronal noise is likely to scale relative to the activity norm. For these reasons, during the optimization of $\mathbf{u}$ for noise robustness, we scaled the variance of the initial noise by the average squared norm of the rates during motif production (see section 4.2), with the aim of ensuring that the optimization would not cause a large increase the activity norm.

Here, in Supplementary Fig. S.3f, we show the success of our approach as, after optimizing $\mathbf{u}$, the time-average norm of the neuronal activities has a similar magnitude as in the case of a half random loop or in the case of control matrices with little or no non normal amplification. Note that the activity norms can occasionally become very large in the case of random $\mathbf{u}$ or control matrices, as the readout and the eigenvector of interest may be orthogonal by chance. In contrast, the activity norms stay well-bounded in the case of tuned $\mathbf{u}$.

###### 5.2.5 Synaptic weight distributions in our thalamocortical model

In the main text, we always display the probability distribution functions (pdfs) of the effective perturbation weights that are contained within the matrix that gets added to the corticocortical matrix $\mathbf{J}_{\text{cc}}$, i.e.:

- during motif production, a motif-specific rank-one matrix: $\mathbf{u}\mathbf{v}^\intercal$, where $\mathbf{u}$ and $\mathbf{v}$ are respectively a thalamocortical and corticothalamic weight vector of a single loop, and
- during the motif preparation period, the matrix $\mathbf{J}_{\text{ct}}\mathbf{S}_{\text{prep}}\mathbf{J}_{\text{tc}}$, where $\mathbf{J}_{\text{ct}}$ is a thalamocortical connectivity matrix and $\mathbf{J}_{\text{tc}}$ is a corticothalamic connectivity matrix.

Therefore, the perturbation matrices have entries that are either a simple multiplication between two individual units' weights in the first case, or a multiplication followed by a sum over the different loops involved in the second case.

Here, we also consider the corresponding distributions of individual units' weights that combine to shape the effective perturbation of $\mathbf{J}_{\text{cc}}$. When separating the effective perturbation weights into individual units' synaptic weights, for each individual corticothalamocortical loop involved, one is free to choose the relative strength of the corticothalamic *vs.* thalamocortical weights. Indeed, if the vectors $\mathbf{u}_0$ and $\mathbf{v}_0^\intercal$ form the effective loop weights $\mathbf{u}_0\mathbf{v}_0^\intercal$, then we can choose scale factors $s_\mathbf{u}$ and $s_\mathbf{v}$ such that $s_\mathbf{u}s_\mathbf{v} = 1$ and the vectors $\mathbf{u} = \frac{\mathbf{u}_0}{s_\mathbf{u}}$ and $\mathbf{v}^\intercal = \frac{\mathbf{v}_0^\intercal}{s_\mathbf{v}}$ can be combined into the same effective loop $\mathbf{u}\mathbf{v}^\intercal = \mathbf{u}_0\mathbf{v}_0^\intercal$. Here, we choose to evenly break the strength of the effective perturbation in between the two sides of the loop, by setting $s_\mathbf{u} = |\mathbf{u}_0| / \sqrt{|\mathbf{u}_0|\,|\mathbf{v}_0|}$ and $s_\mathbf{v} = |\mathbf{v}_0| / \sqrt{|\mathbf{u}_0|\,|\mathbf{v}_0|}$.

The resulting full distributions of individual loop units' weights are shown in Supp. Fig. S.4b,d,f,h,j; along with the full distributions of effective perturbation weights for comparison (Supp. Fig. S.4a,c,e,g,i). To better visualize the shape and the relative scale of the loop weights, we also plot the $\mathbf{J}_{\text{cc}}$ weights (which are Gaussian by construction), as well as a Gaussian distribution with parameters that are matched to the loop weights.

We note that in the case of motif production weights with a half-random perturbation (i.e., one of the loop's weight vector is tuned to control a few eigenvalues while the other is Gaussian i.i.d.), the individual units' weights also appear to be Gaussianly distributed (see ref. [81] for a related result; Supplementary Fig. S.4b,f). In contrast, the effective perturbation loop weight for motif production are then distributed as the product of two Gaussians (Fig. 2c,f; Supplementary Fig. S.4a,e).

Further, in the case of fully tuned loops (either one loop for motif production or several loops for motif preparation), the individual units' weights still appear to approximately follow a Gaussian distribution altogether (Supplementary Fig. S.4d,h,j). Interestingly, this result is not trivially imposed through the optimization algorithm (CMA-ES optimizes weights non-locally following a multivariate Gaussian model, but that does not imply that the mixture distribution of all loop weights should be Gaussian).

Despite the fact that the full optimization of the thalamocortical loop weights involved in motif production preserves the shape of the weight distribution compared to a half-random loop, the former optimization still leads to a significant reduction of the magnitude of the weights (Supplementary Fig. S.4 c & d and g & h *vs.* a & b and e & f respectively; Fig. 2 f *vs.* c).

This emphasizes how the full weight optimization we developped leads to reasonably shaped distribution weights, with reasonable weight magnitudes.

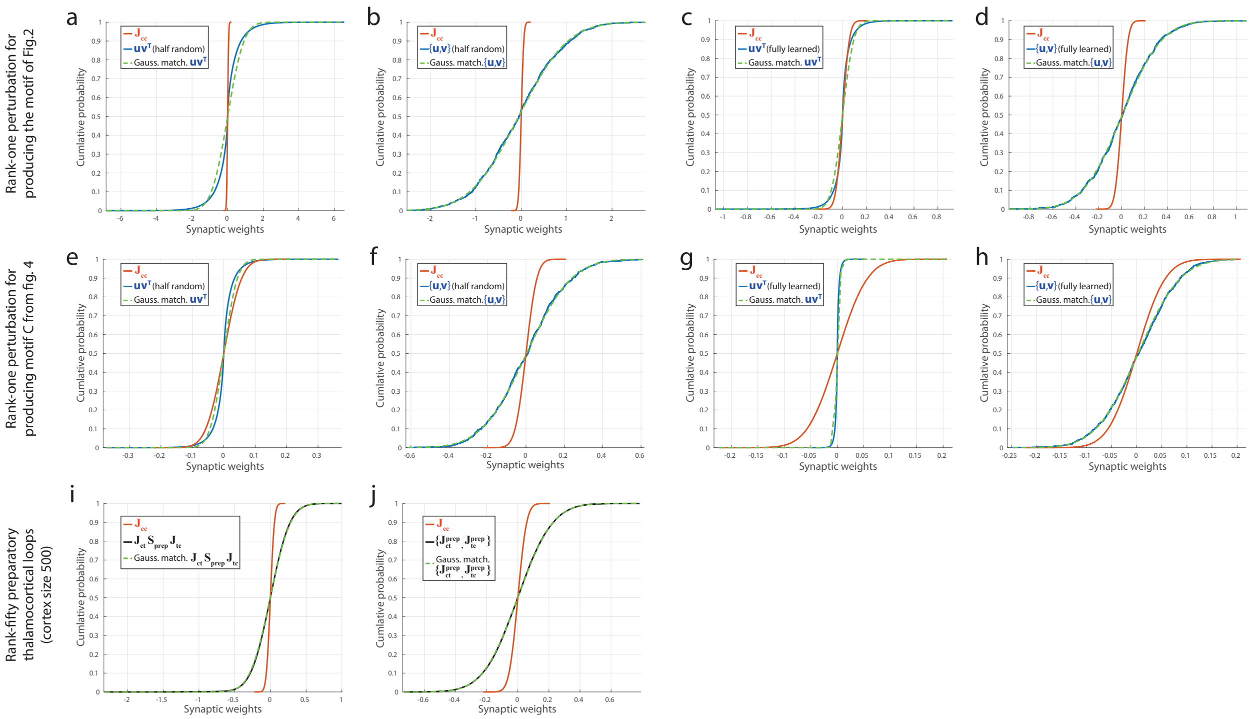


Figure S.4: Cumulative distribution functions (cdfs) for the thalamocortical networks' weights, during *(i)* the production of the sinc function as in Fig. 2 (panels **a**–**d**), *(ii)* the production of motif C as in Fig. 4 (panels **e**–**h**), and *(iii)* the preparatory network as in Fig. 3c–e (panels **i** & **j**). As a reference, the cdf for the weights in $\mathbf{J}_{\rm cc}$ is shown in red in all panels (i.e., a particular drawing from a centered Gaussian distribution with standard deviation $1/\sqrt{N}$ where $N = 500$ is the cortical network's size). In panels (**a**–**h**), the blue curve is the cdf for the weights of a rank-one perturbation tuned for motif production: either half-tuned weights which set the target eigenvalues (panels **a** & **b** and **e** & **f**) or fully-tuned weights which are also optimized for noise robustness (panels **c** & **d** and **g** & **h**). In addition, we show both the effective perturbation weights (i.e., the entries of $\mathbf{u}\mathbf{v}^\intercal$; panels **a**, **c**, **e** and **g**) or the individual weights between units (i.e., the mixture distribution gathering the entries of $\mathbf{u}$ and $\mathbf{v}$, symbolized by $\{\mathbf{u}, \mathbf{v}\}$; panels **b**, **d**, **f** and **h**). Finally, in panels **i** & **j**, we plot in black the cdf of weights to and from a 50-neurons preparatory thalamic population, either in the form of the entries of the effective perturbation matrix $\mathbf{J}_{\rm ct}\mathbf{S}_{\rm prep}\mathbf{J}_{\rm tc}$ (panel **i**) or in the form of the individual weights between units (i.e., the mixture distribution gathering the individual thalamocortical and thalamocortical weights separately for each loop and symbolized as $\{\mathbf{J}_{\rm ct}^{\rm prep}, \mathbf{J}_{\rm tc}^{\rm prep}\}$, panel **j**). In all panels, we also plot a Gaussian cdf with mean and variance matched to the thalamocortical loop distributions for comparison (dashed green line).

#### 5.3 Scaling of the neural resources as a function of motif complexity and sequence length

Here, we summarize how the capacity of our network scales with number of units. For our task, the notion of capacity actually encompasses two different aspects: first, the complexity of the motifs that the network can approximate well – which can be mathematically quantified as the number and the eccentricity of the eigenvalues required to match a desired motif (see Supplementary Fig. S.2) – and, second, the number of motifs that the network can store.

As we illustrate in Supplementary Fig. S.5a, in our model, we have shown that sequences of motifs can be generated via the selective disinhibition of thalamic subpopulations with specific weights to and from cortex that control the cortical dynamics. Each motif therefore requires a dedicated thalamic unit (or set of units) to generate the eigenmodes from which it is constructed. Thus, it is clear that to generate a sequence of length $N_{\rm motifs}$ motifs – where each motif is constructed from its own set of eigenmodes – the number of thalamic units required scales with $N_{\mathrm{motifs}}$. The cortical population, on the other hand, scales with the number of eigenmodes needed to generate a single motif ($N_{\mathrm{modes/motif}}$; see Supplementary Figs. S.1 & S.2).

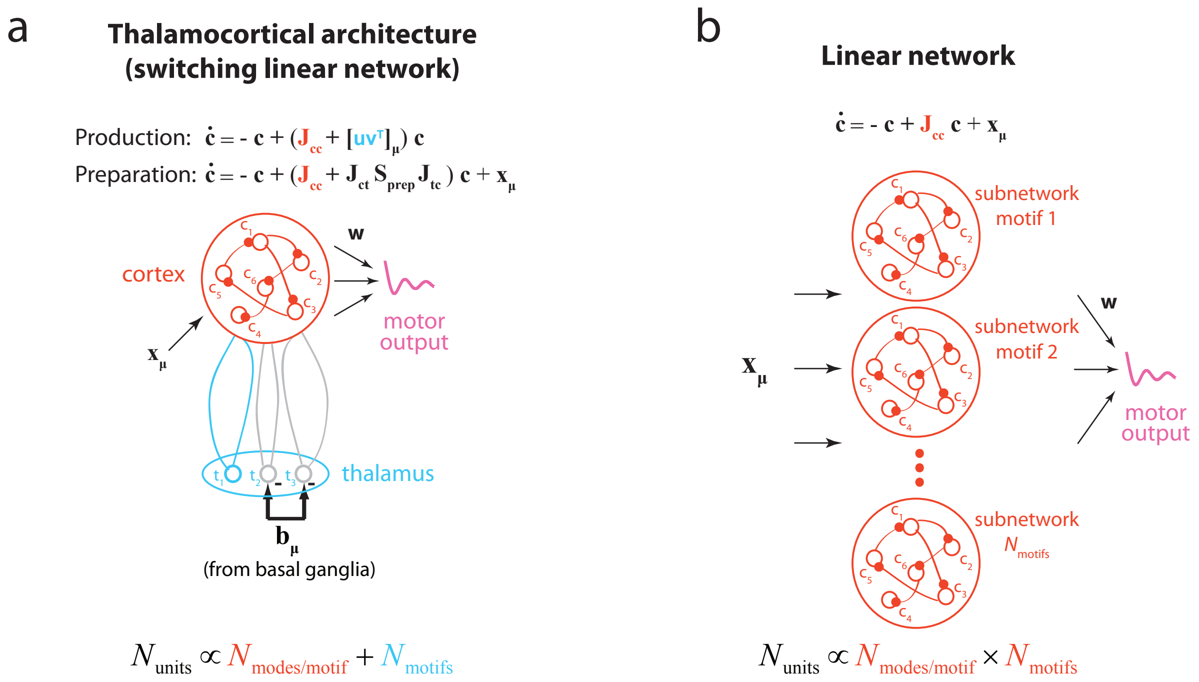


Figure S.5: Scaling of the number of units needed to produce sequences of length $N_{\mathrm{motifs}}$ assuming the number of eigenmodes needed to approximate a typical motif is $N_{\mathrm{modes/motif}}$ (see Fig. S.2). **a**) The corticothalamocortical architecture. By including a rectifying non-linearity in thalamus, the network behaves as a switching linear system where different motifs are encoded separately through the parameters corresponding to specific corticothalamocortical loops. These loops remap the cortical dynamics to generate the desired motif-specific outputs, and thus permit the reuse of the cortical units for each motif. Hence, the total number of units in the network scales as $N_{\mathrm{motifs}} + N_{\mathrm{modes/motif}}$. **b**) A linear system. The production of more complex sequences is possible via motif-specific inputs that instruct the network's output. However, these inputs cannot remap the underlying dynamics, and so the number of eigenmodes that can be generated scales with the number of units in the network. Thus, a sequence of length $N_{\mathrm{motifs}}$ requires a network of size $N_{\mathrm{motifs}} \times N_{\mathrm{modes/motif}}$. This can be visualized as a collection of subnetworks, one for each motif.

As a sanity check, it is interesting to compare the capacity of our thalamocortical network to the capacity of a matched linear network (Supplementary Fig. S.5). Our network possesses a rectifying non-linearity in thalamus, and therefore it has the potential to achieve a superior capacity. Note however that, even if nonlinear dynamics always has the potential to achieve more powerful computations than linear dynamics, this potential is not necessarily achieved, notably if the network structure and/or weight adjustments are not well-adapted to the task. For instance, nonlinear networks cannot be trivially trained with gradient descent to robustly perform flexible and extensible motor sequencing in a similar way as our model does [31]. In addition, a non-linear model in which all plastic synapses are adjusted when learning a motif will suffer from learning interference. Therefore, more complex models do not necessarily perform better than a simple, but well-designed, model.

In order to perform our flexible sequencing task (Fig. 1a), a linear model would require a number of units proportional to the total number of eigenvalues needed by all motifs in a sequence, or $N_{\mathrm{motifs}} \times N_{\mathrm{modes/motif}}$ for complex motifs that are each constructed from different timescales (Supplementary Fig. S.5b). This number grows considerably faster with the number of motifs than the total number of units in our model (Supplementary Fig. S.5a).

Hence, our model is able to leverage the larger expressivity of nonlinear dynamics. This stems from the interplay between the thalamocortical network's architecture and the nonlinearity, which allows different thalamic units to act as powerful controllers of cortical dynamics in correspondance to the task's structure. Notably, the absence of excitatory recurrence in thalamus *(i)* creates an opportunity for faster responses [83, 72] which are key for its ability to shape the slower recurrent cortical dynamics during each motif [63] and *(ii)* helps countering interference during the learning and execution of different motifs.

##### 5.4 Introducing a plausible thalamic timescale and rectified dynamics without losing theoretical insights

Here, we use the methodology we developed in Supplementary Sec. 5.1.5 to show that two main simplifications we made relative to biological neuronal dynamics may be relaxed while making a minimal impact on the dynamics, and hence without losing the insights from the theoretical analysis that we present in the main text.

More specifically, we explain in Supplementary Sec. 5.1.5 how the deviations induced by the presence of a thalamic timescale can be treated as a continuous biased noise whose effect can be minimized through the adjustment of the vector $\mathbf{u}$ while eigenvalue control is performed as in the main text. In Supplementary Fig. S.6a, we demonstrate that taking a thalamic timescale that is just 10 times faster than the cortical timescale (an order of magnitude compatible with the difference between single-neuron timescale and recurrent population dynamics timescale [83, 72]) induces very small deviations (grey curves) compared to the case of instantaneous thalamus (black curves) when starting motifs from their ideal initial rate patterns (Eq. 5). Also, the finite response timescale of thalamus does not prevent fast convergence to appropriate initial conditions during the motifs' preparatory period using as small of a network as in Fig. 3 (10% of the cortical network with connectivities devised as in Supplementary Sec. 5.1.5; Supplementary Fig. S.6b, green curve).

Finally, in Supplementary Sec. 5.1.5, we also show how to design a constant bias to the dynamics such that non-rectified rates would only hit negative values rarely. We show in Supplementary Fig. S.6a that, then, introducing a rectification to the dynamics (the cyan curve, following Eq. S.23 vs. the green line, following Eq. S.16) leads to minimal deviations during motifs. Interestingly, in Supplementary Fig. S.6b, we also show how introducing the rectifying non-linearity in the dynamics usually speeds the preparatory process (inset).

In conclusion, these results show that our framework for sequence production can successfully be generalized to more biologically constrained positive rectified dynamics with realistic timescale ranges.

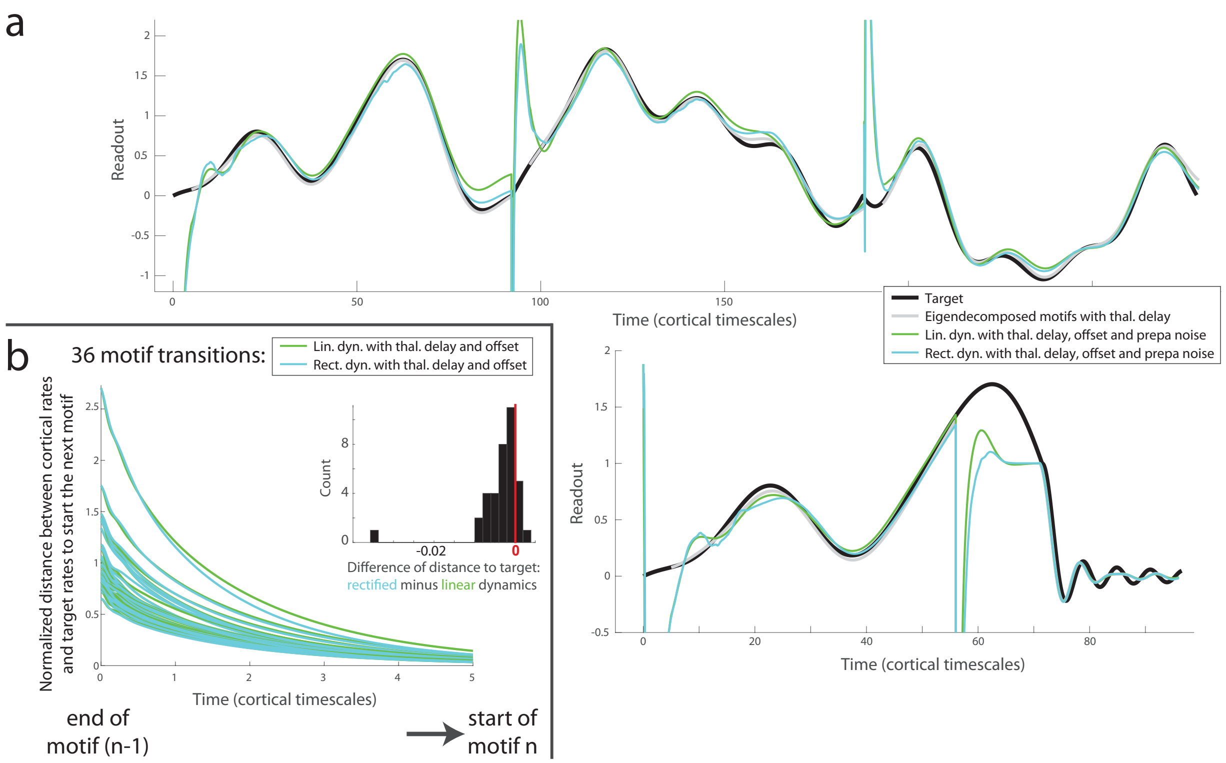


Figure S.6: **a)** The readout for a network trained to produce different sequences, while accounting for important biological constraints, as compared to the desired sequence (black). First, thalamic units are constrained to be realistically fast (10 times faster than the cortical units). The eigenvalues are still constrained using the same theoretical framework as in the main text (Eq. 8) which assumes an idealized network with immediate thalamic responses, but the deviations induced by the thalamic delay are interpreted as a biased noise whose effects on the readout are minimized along with the effect of initial rates noise (Eq. S.19). The effect of introducing the thalamic delay on motif production is visible by comparing the black target readout with the grey lines, which show the cortical readout when exactly adjusting the rates to their ideal values at the beginning of each motif. In addition, the rates are constrained to be mostly positive in the linear regime by adding motif-specific constant inputs to the dynamics (Supplementary Sec. 5.1.5). The green line is the readout of these offset linear dynamics, while the cyan line is the result when additionally rectifying the rates below zero. Note that we used a restricted number of thalamic units to optimize the preparatory network (10% of the cortical size or 50 units). In consequence, for more complex motifs (like the sinc function; inset), a longer preparatory period is needed to make sure that the activities are not too different from the noise-free linear and positive solution, or else the readout of the rectified dynamics will differ from the target (not shown). **b)** Details of the effectiveness of the preparatory network constrained by the presence of a thalamic delay and of constant inputs setting the rates to mostly positive values during linear dynamics. The x-axis is restricted to the start of the preparatory period which links two motifs, and 36 motif transitions are illustrated. Each trace shows the euclidean distance between the cortical rates and the target cortical rate pattern for starting the next motif, relative to the norm of the target cortical pattern. The green lines correspond to linear dynamics while the rectified dynamics are shown in cyan. Inset: the difference of final distance from the target rates between rectified and linear dynamics, showing that the rectified dynamics often cause the cortical rates to converge faster than in the case of linear dynamics.

[S63] Gene Franklin. *Feedback control of dynamic systems*. Pearson, Boston, 2015.

[S64] S. Ganguli, D. Huh, and H. Sompolinsky. Memory traces in dynamical systems. *Proc. Natl. Acad. Sci. U.S.A.*, 105(48):18970–18975, Dec 2008.

[S65] Wulfram Gerstner, Werner M. Kistler, Richard Naud, and Liam Paninski. *Neuronal Dynamics*. Cambridge University Press, 2009.

[S66] Guillaume Hennequin, Tim P. Vogels, and Wulfram Gerstner. Non-normal amplification in random balanced neuronal networks. *Physical Review E*, 86(1), jul 2012.

[S67] Holly P. Hirst and Wade T. Macey. Bounding the roots of polynomials. *The College Mathematics Journal*, 28(4):292, September 1997.

[S68] Xin Jin and Rui M. Costa. Start/stop signals emerge in nigrostriatal circuits during sequence learning. *Nature*, 466(7305):457–462, jul 2010.

[S69] Xin Jin, Fatuel Tecuapetla, and Rui M. Costa. Basal ganglia subcircuits distinctively encode the parsing and concatenation of action sequences. *Nat Neurosci*, 17(3):423–430, Mar 2014.

[S70] T. Kaneko. Local connections of excitatory neurons in motor-associated cortical areas of the rat. *Front Neural Circuits*, 7:75, 2013.

[S71] Jeongjin Kim, Youngsoo Kim, Ryuichi Nakajima, Anna Shin, Minju Jeong, Ah Hyung Park, Yongcheol Jeong, Seonmi Jo, Seungkyoung Yang, Hosung Park, Sung-Hwan Cho, Kwang-Hyun Cho, Insop Shim, Jae Hoon Chung, Se-Bum Paik, George J. Augustine, and Daesoo Kim. Inhibitory basal ganglia inputs induce excitatory motor signals in the thalamus. *Neuron*, 95(5):1181–1196.e8, August 2017.

[S72] Sukbin Lim and Mark S. Goldman. Balanced cortical microcircuitry for maintaining information in working memory. *Nat Neurosci*, 16(9):1306–1314, Sep 2013.

[S73] Laureline Logiaco, Laurence Abbott, and Gary S. Escola. A thalamocortical model solves temporal sequencing tasks more robustly than rnns trained with stochastic gradient descent. In *Computational and Systems Neuroscience*, 2019.

[S74] Dhagash Mehta, Tianran Chen, Tingting Tang, and Jonathan D. Hauenstein. The loss surface of deep linear networks viewed through the algebraic geometry lens. *arXiv e-prints*, page arXiv:1810.07716, Oct 2018.

[S75] Skander Mensi, Richard Naud, Christian Pozzorini, Michael Avermann, Carl C. H. Petersen, and Wulfram Gerstner. Parameter extraction and classification of three cortical neuron types reveals two distinct adaptation mechanisms. *Journal of Neurophysiology*, 107(6):1756–1775, March 2012.

[S76] Francesco Mezzadri. How to generate random matrices from the classical compact groups. *arXiv e-prints*, pages math–ph/0609050, Sep 2006.

[S77] B. K. Murphy and K. D. Miller. Balanced amplification: a new mechanism of selective amplification of neural activity patterns. *Neuron*, 61(4):635–648, Feb 2009.

[S78] Srdjan Ostojic and Nicolas Brunel. From spiking neuron models to linear-nonlinear models. *PLoS Computational Biology*, 7(1):e1001056, jan 2011.

[S79] Alfonso Renart, Jaime de la Rocha, Peter Bartho, Liad Hollender, Néstor Parga, Alex Reyes, and Kenneth D. Harris. The asynchronous state in cortical circuits. *Science*, 327(5965):587–590, Jan 2010.

[S80] Abigail A. Russo, Sean R. Bittner, Sean M. Perkins, Jeffrey S. Seely, Brian M. London, Antonio H. Lara, Andrew Miri, Najja J. Marshall, Adam Kohn, Thomas M. Jessell, Laurence F. Abbott, John P. Cunningham, and Mark M. Churchland. Motor cortex embeds muscle-like commands in an untangled population response. *Neuron*, 97(4):953–966.e8, feb 2018.

[S81] Friedrich Schuessler, Alexis Dubreuil, Francesca Mastrogiuseppe, Srdjan Ostojic, and Omri Barak. Dynamics of random recurrent networks with correlated low-rank structure. *arXiv e-prints*, page arXiv:1909.04358, Sep 2019.

[S82] Bettina C. Schwab, Daisuke Kase, Andrew Zimnik, Robert Rosenbaum, Jonathan E. Rubin, and Robert S. Turner. Weak modulation of thalamic discharge by basal ganglia output in association with a reaching task. February 2019.

[S83] H.Sebastian Seung, Daniel D. Lee, Ben Y. Reis, and David W. Tank. Stability of the memory of eye position in a recurrent network of conductance-based model neurons. *Neuron*, 26(1):259–271, apr 2000.

[S84] K. V. Shenoy, M. Sahani, and M. M. Churchland. Cortical control of arm movements: a dynamical systems perspective. *Annu. Rev. Neurosci.*, 36:337–359, Jul 2013.

[S85] S. M. Sherman. Tonic and burst firing: dual modes of thalamocortical relay. *Trends Neurosci.*, 24(2):122–126, Feb 2001.

[S86] S Murray Sherman. Thalamus plays a central role in ongoing cortical functioning. *Nature Neuroscience*, 19(4):533–541, apr 2016.

[S87] D. Sussillo, M. M. Churchland, M. T. Kaufman, and K. V. Shenoy. A neural network that finds a naturalistic solution for the production of muscle activity. *Nat. Neurosci.*, 18(7):1025–1033, Jul 2015.

[S88] Terence Tao. Outliers in the spectrum of iid matrices with bounded rank perturbations. *Probability Theory and Related Fields*, 155(1-2):231–263, nov 2011.

[S89] C. van Vreeswijk and H. Sompolinsky. Chaos in neuronal networks with balanced excitatory and inhibitory activity. *Science*, 274(5293):1724–1726, Dec 1996.

[S90] C. van Vreeswijk and H. Sompolinsky. Chaotic balanced state in a model of cortical circuits. *Neural Comput*, 10(6):1321–1371, Aug 1998.
